# Identification of Parkinson’s disease-associated regulatory variants in human dopaminergic neurons reveals modulators of *SCARB2* and *BAG3* expression

**DOI:** 10.64898/2026.03.26.714241

**Authors:** Deborah Gérard, Jochen Ohnmacht, Borja Gomez-Ramos, Marie Catillon, Jafar Sharif, Nina Baumgarten, Dennis Hecker, Aurelien Ginolhac, Zied Landoulsi, Elena Valceschini, Aleksandar Rakovic, Christine Klein, Patrick May, Haruhiko Koseki, Marcel H. Schulz, Thomas Sauter, Rejko Krüger, Lasse Sinkkonen

**Affiliations:** Department of Health, Medicine and Life Sciences (DHML), University of Luxembourg, L-4365 Belvaux, Luxembourg; Translational Neuroscience, Luxembourg Centre for Systems Biomedicine (LCSB), University of Luxembourg, L-4365 Belvaux, Luxembourg; Laboratory for Developmental Genetics, RIKEN Center for Integrative Medical Sciences (IMS), Yokohama, 230-0045, Japan; German Centre for Cardiovascular Research, Partner site Rhein-Main, D-60590 Frankfurt am Main, Germany; Cardio-Pulmonary Institute, Goethe University, D-60590, Frankfurt am Main, Germany; Institute for Computational Genomic Medicine, Goethe University, D-60590 Frankfurt am Main, Germany; Transversal Translational Medicine, Luxembourg Institute of Health (LIH), L-1445, Luxembourg, Luxembourg; Institute of Neurogenetics, University of Lübeck, D-23538, Lübeck, Germany; Department of Cellular and Molecular Medicine, Graduate School of Medicine, Chiba University, Chiba 260-8670, Japan; Parkinson Research Clinic, Centre Hospitalier de Luxembourg (CHL), L-1210, Luxembourg, Luxembourg

**Keywords:** Dopaminergic neurons, Parkinson’s disease, regulatory variants, enhancers, transcription factors

## Abstract

A hallmark of Parkinson’s disease (PD) is the degeneration of midbrain dopaminergic neurons (mDANs). Genome-wide association studies (GWAS) have identified single nucleotide polymorphisms (SNPs) associated with PD, but causal variants and mechanisms remain unknown. Many PD-associated SNPs reside in regulatory regions, where they may disrupt transcription factor binding sites (TFBS) and alter gene expression. To assess how non-coding PD SNPs affect gene regulation in mDANs, we identify variants predicted to alter TF binding and functionally validate their effects in a cell type-specific context. We integrate time-series transcriptome and chromatin accessibility data from iPSC-derived neurons with chromatin topology and genetic variants. We profile 3D chromatin conformation in neuronal progenitors (smNPCs) and mDANs using LowC, identifying changes in A/B compartments and topologically associated domains. PD SNPs are enriched near genes expressed in mDANs, and we predict 254 regulatory variants that create or disrupt TFBS. Using chromatin conformation data, we link variants to target genes. At the BAG3 and SCARB2 loci, reporter assays in mDANs show reduced transcription driven by PD-associated alleles. Knock-down of NR2C2, a putative SCARB2 regulator, increases SCARB2 expression in differentiating neurons. The PD-associated SCARB2 allele shows reduced chromatin accessibility in mDANs and is associated with decreased expression in brain eQTL data. Insertion of PD-associated BAG3 allele by prime editing reduces chromatin accessibility across cell types, consistent with altered binding of LIM-homeodomain transcription factors. Together, these results prioritize functional PD SNPs and show that variants at SCARB2 and BAG3 modulate gene expression in mDANs, providing mechanistic insight into PD.

## Introduction

Parkinson’s disease (PD) is the fastest growing neurodegenerative disorder, affecting millions worldwide with numbers expected to double within the next two decades (1). Its hallmark is the selective degeneration of midbrain dopaminergic neurons (mDANs), which leads to the well-documented motor symptoms of PD (2). While the etiology of PD is complex, involving both genetic and environmental factors, genetic predispositions play a crucial role in the disease’s development and progression (3).

Large-scale genomic projects, such as the 1000 Genomes Project, have catalogued millions of genetic variants across diverse human populations, and together with efforts to map the human epigenome, such as ENCODE and IHEC, have revealed that the vast majority of these variants are non-coding (4–7). These non-coding variants are central in regulating gene expression, particularly through transcription factor binding sites (TFBS) located within enhancers and promoters, and can influence complex traits and diseases primarily by modulating the activity of these regulatory regions (8,9).

The effects of non-coding variants are largely cell type-specific, depending on the enhancers that are active in a particular cell type. For instance, a variant located within an enhancer active in mDANs might have no effect in other cell types, such as liver or muscle cells, where that enhancer is not active. This specificity arises as each cell type has a unique set of transcription factors that are expressed and capable of binding to accessible regions within the genome. For example, the non-coding variants associated with Alzheimer’s disease are typically enriched in enhancers active in microglia (10), often leading to more severe immune response to the accumulation of tau fibrils during disease progression (11).

Expression quantitative trait loci (eQTL) analysis has shown that the influence of *cis*-regulatory variants on gene expression often manifests as subtle changes, typically altering gene expression only by 10% to 50%. Despite these modest alterations, the cell type-specific impact can be profound, affecting disease susceptibility and progression as *trans*-regulatory variants through a cascade of regulatory events (12). Furthermore, the regulatory effects of non-coding variants are mediated through networks of enhancers and transcription factors, each of which may only be active under certain physiological conditions or stages of cell differentiation (13). This nuanced regulatory landscape underscores the necessity of studying these variants in cell type-specific models to fully understand their roles in disease pathogenesis (14,15). For example, expression of the SNCA gene, which encodes alpha-synuclein, a key protein in PD pathology, has been found to be under control of neuron-specific enhancers carrying non-coding variants associated with disease risk (16).

In the context of PD, genome-wide association studies (GWAS) have identified numerous single nucleotide polymorphisms (SNPs) associated with the disease, most of which are located in non-coding regions (17). These variants are expected to play an important role in explaining the missing heritability of PD. Cumulative small effects in gene regulatory processes - resulting in expression changes in genes playing crucial roles in PD pathogenesis – can modify PD susceptibility. However, these interactions have not yet been mapped in detail for PD nor many other complex disease (18).

Non-coding variants associated with PD are particularly observed in the proximity of genes associated with cellular functions such as mitochondrial or lysosomal homeostasis (19). Dysfunctions in these pathways are a notable feature in PD, and PD-associated variants are often located near genes involved in these pathways, suggesting a potential regulatory impact by these SNPs on lysosomal and other crucial pathways.

Here, we have utilized a novel integrative approach combining time-series transcriptomic, chromatin accessibility, and chromosome conformation data from induced pluripotent stem cell (iPSC)-derived mDANs to explore how PD-associated non-coding variants can affect gene regulation. Our analysis focused on identifying PD SNPs that can alter TF binding in cell type-specific regulatory regions. This led to the identification of hundreds of candidate variants for detailed functional studies. From these rs1465922 at the *SCARB2* locus and rs144814361 at the *BAG3* locus were found to create new TF binding sites for NR2C2 and LIM homeodomain (HD-LIM) TFs such as LHX1, respectively. These variants modulate SCARB2 and BAG3 expression and, for SCARB2, were linked to lower expression in critical brain regions like the substantia nigra. These findings advance our understanding of how non-coding variants contribute to PD pathogenesis and highlight some molecular mechanisms that can drive disease susceptibility and inform future disease-modifying treatments.

## Materials and Methods

### Cell lines

The human iPSC line GM17602 (Coriell) was used in this study as a control for differentiation and for the generation of a tyrosine hydroxylase (TH) reporter cell line (TH-Rep1). This iPSC line was previously described in (20) and used in (21) for the generation of the reporter line as already outlined (22). The second reporter iPSC line (TH-Rep2) was generated similarly using STBCi033-B cell line (22). Cell lines were regularly tested for mycoplasma.

### Cell culture and differentiation

The TH-Rep1 cell line was used during all experiments. Maintenance of small molecule neural precursor cells (smNPC) and differentiation towards mDANs are described in (23). smNPC were grown in N2B27 medium constituted of 50% of Dulbecco’s Modified Eagle Medium:Nutrient Mixture F-12 (DMEM/F-12) media (Gibco, ThermoFisher Scientific, 21331-046), 50% of NeurobasalTM medium (Gibco, ThermoFisher Scientific, 21103-049), 0.5% N-2 Supplement (100X) (Gibco, ThermoFisher Scientific, 17502-001), 1% B-27TM Supplement (50X) (minus vitamin A) (Gibco, ThermoFisher Scientific, 12587-001), 1% Penicillin-Streptomycin (10000 u/mL) (Gibco, ThermoFisher Scientific, 15140-122), GlutaMaxTM Supplement (Gibco, ThermoFisher Scientific, 35050-061) supplemented with 500 nM Purmorphamine (PMA) (Sigma-Aldrich, SML0868), 150 µM L-Ascorbic acid (AA) (Sigma-Aldrich, A4544) and 3 µM CHIR99021 (CHIR) (Axon Medchem, AXON1386) in a constant atmosphere of 37°C and 5% CO2. All experiments were performed with cells passaged for less than 40 times (23). Neuronal differentiation was initiated on day 0 by adding differentiation media 1 (DIFF1) (N2B27 supplemented with 1 µM PMA Sigma-Aldrich, SML0868, 200 µM AA (Sigma-Aldrich, A4544)) and 100 ng/mL of Fibroblast Growth Factor-8 (FGF8b) (Peprotech, 100-25)). From day 8 on, cells were split and plated on fresh Geltrex (Gibco, A1413302) and differentiation media 2 (DIFF2) (N2B27 supplemented with 0.5 µM PMA (Sigma-Aldrich, SML0868) and 200 µM AA (Sigma-Aldrich, A4544))) was added to the cells. From day 10 on, maturation media (N2B27 supplemented with 200 µM AA (Sigma-Aldrich, A4544)), 500 µM dibutyryl cAMP (dbcAMP) (Santa Cruz, sc-201567C), 10 ng/mL brain derived neurotrophic factor (BDNF) (Peprotech, 450-02-100UG), 10 ng/mL glial derived neutrophic factor (GDNF) (Peprotech, 450-10-1MG) and 1 ng/mL transforming growth factor-β3 (TGFB3) (Peprotech, 100-36E-10UG). Media was replaced every two days.

### Bacterial culture, plasmid extraction and lentivirus production

Bacteria harbouring the target plasmid were retrieved from glycerol stocks, transferred to a skirt tube (Greiner bio-one, 187262) containing 5 ml of LB Broth medium (Sigma-Aldrich, L7658) supplemented with 100 µg/ml ampicillin (Sigma-Aldrich, A9518-5G). After incubation at 37°C and shaking at 120 rpm for 5 hours the bacterial culture was inoculated into an Erlenmeyer flask containing 150 mL of LB Broth medium supplemented with 100 µg/ml ampicillin, followed by an overnight incubation at 37°C with agitation at 120 rpm. After bacterial expansion, plasmid extraction was performed using NucleoBond Xtra Midi EF (Macherey-Nagel, 740420.50) according to the manufacturer’s instructions.

Lentiviral particles were produced by transfecting HEK-293T cells with the desired plasmids. 8 million HEK293T cells were initially seeded in 15 ml of DMEM (Gibco, ThermoFisher Scientific, 41965062) supplemented with 1% Pen/Strep (Gibco, ThermoFisher Scientific, 15140-122) and 10% fetal bovine serum (FBS, ThermoFisher Scientific, 26400044) in a T75 flask. Third-generation lentiviral particles were generated by mixing 4 µg of pMDG, 2 µg of pMDL, 2 µg of pREV, and 8 µg of the desired plasmid with 200 µL of CaCl_2_ (Sigma-Aldrich, 21115-100ML) and sterile water was added to reach a volume of 800 µL. This 800 µL of plasmid mixture was slowly bubbled into 800 µL of HEPES buffered saline (Sigma-Aldrich, 51558-50ML) in a dropwise manner and incubated for 20 min at room temperature. Meanwhile, 16 µL of 25 mM chloroquine (Sigma-Aldrich, C6628) was added to the T75 flask containing the HEK-293T cells and incubated for a minimum of 5 min to aid transfection. Then, the transfection mixture was added to the HEK-293T cells. After 6 hours, the medium was replaced with 14 mL of fresh medium. After 48 hours, lentiviral particles were concentrated using a sucrose gradient. Briefly, the supernatant of HEK293T cells containing the newly produced lentiviral particles was overlayed with a concentrator solution (10% sucrose w/v (Merck, 1.07687.1000), Tris-HCl (VWR, A1086.1000) pH 7.4 50 mM, NaCl (Carl Roth, 3957.1) 100 mM, Ethylenediamine tetraacetic acid (EDTA) (Carl Roth, CN06.3) 0.5 mM) at a 4:1 v/v ratio and centrifuge at 9500x*g* for 4 hours at 4°C. Following the centrifugation step, the supernatant was discarded and the pellets containing the lentiviral particles of interest were resuspended into Hank’s balanced salt solution (HBSS) (Gibco, ThermoFisher Scientific, 14025092) and incubated overnight at 4°C. The next day, the lentiviral particles were aliquoted and stored at −80°C until further use. To evaluate transduction efficiency and track GFP expression levels via flow cytometry, a standard curve was generated using varying volumes of lentiviral particles.

### Transduction of shRNAs and luciferase vectors

Before transduction, 8 days differentiated neurons were split and 300 000 neurons were seeded in DIFF2 media in freshly coated 12 well plates. The next day, 80 µL of concentrated lentiviral particles were diluted to 420 µL of DIFF2 media and this mixture was added to neurons followed by a spinoculation at 250x*g* for 10 minutes at 21°C. After 6 hours, the media was replaced with fresh DIFF2 media. Cells transduced with shRNAs against *LHX1* and *NR2C2* were collected 72 hours post-transduction and 6 days post-transduction for cells transduced with shRNAs against *NR2C2.* Transduction efficiency was assessed by quantifying GFP expression levels by flow cytometry.

Supernatant of cells transduced with the luciferase vectors were collected 48 hours post-transduction and luciferase levels were measured using the Secreted-Pair Dual Luminescence assay kit (GeneCopoeia, LF032) according to the manufacturer’s instructions.

### Generation of TH-Rep1-SNP-BAG3 cell line

#### Precise SNP-BAG3 editing in human induced pluripotent stem cells (iPSCs) using the Prime Editing system

After two passages post-thawing, iPSCs TH-Rep1 maintained in mTeSR™ Plus medium (StemCell^TM^ Technologies, 100-0276) were nucleofected with a mix of 3 plasmids containing pCMV-PE2, pU6-pegRNA_SNP and pE3b-BAG3, or pmaxGFP^TM^ control vector (Lonza), using the 4D-Nucleofector TM Core + X Unit (Lonza) instrument and the P3 Primary Cell Kit (Lonza V4XP-3024) (program CA137). Cells were immediately seeded in Matrigel-coated plates in the presence of 10 µM ROCK inhibitor to promote survival and attachment. Following recovery and expansion, clonal isolation was performed at 70% confluency by limiting dilution into 96-well plates. Individual colonies were screened by TaqMan™ genotyping and confirmed by Sanger sequencing. The stemness markers of edited iPSC TH-Rep1-SNP-BAG3 clone (1/34) was assessed as described below by RTqPCR and immunocytochemistry for OCT3/4, NANOG, and DNMT3B. pCMV-PE2 (Addgene plasmid # 132775) and pU6-pegRNA-GG-acceptor (Addgene plasmid # 132777) were a gift from David Liu. BPK1520 was a gift from Keith Joung (Addgene plasmid # 65777).

The SNP variation rs144814361 (C/T) was inserted into iPSC TH-Rep1 genome at the position BAG3 GRCh38 Ch10:119651405 by prime editing. The pegRNA sequence containing the SNP variant was designed by using the pegFinder tool (http://pegfinder.sidichenlab.org) (24).

The components sgRNA, scaffold and 3’ extension were annealed to generate the pegRNA fragment and cloned into the plasmid pU6-pegRNA-GG-acceptor by NEBridge^©^Golden Gate assembly reaction using the enzyme BsaI-HF^®^ v2 (NEB E1601S). Each component resulted in the annealing of specific oligos: sgRNA (sgF_sgR), scaffold (scaffF_scaffR) and 3’ extension (extensF_extensR). See the below table of oligonucleotides for sequence details.

The designed sequence of full-length pegRNA was CTGGTCCAGTCCAGAGAAGTGTTTTAGAGCTAGAAATAGCAAGTTAAAATAAGGCTAGTCCGTTATCAACTTGAAAAAGTGGCACCGAGTCGGTGCGGCCGCGGCCAATTTCTCTGGAC TGGA.

PE3b nick sgRNA was designed to target and nick specifically the non-edited DNA strand. The fragments PE3b_sgF and PE3b_sgR were annealed and cloned into BPK1520 by NEBridge^©^Golden Gate assembly reaction using the enzyme BsmBI-v2 (NEB R0739S).

### Neural induction of iPSC TH-Rep1-SNP-BAG3 in Neural Progenitor Cells (NPCs)

According to the protocol STEMdiff™ Neural System (StemCell), iPSCs were seeded at 2×10^6^ cells/well (6 well-plate) in STEMdiff™ SMADi Neural Induction medium (StemCell, cat 08581) and 10 µM ROCK inhibitor. Medium was changed daily for 14 days, and cells were passed as necessary (1:6 to 1:10) to maintain optimal density. After 20 days, cells were seeded at 4×10^5^ cells/well (6 well-plate) in smNPC medium.

The absence of stemness and appearance neural progenitor markers during neural induction of iPSC into NPC TH-Rep1-SNP-BAG3 was assessed by RTqPCR as described below, using stemness markers OCT3/4, NANOG, and DNMT3B and neural progenitor markers NESTIN, SOX1 and PAX6.

### Characterization of TH-Rep1-SNP-BAG3 cell line

#### TaqMan Genotyping

The presence of the BAG3 SNP variant was assessed in 34 iPSC clones using TaqMan® SNP Genotyping Assays human (Thermofisher, Cat. No. 4351379, Assay ID: C_160612363_10, SNP ID: rs144814361). One clone REP1-SNP-BAG3-TH-mcherry (named TH-Rep1-SNP-BAG3) was herezygous for SNP BAG3 (C/T).

#### Sanger sequencing

Genomic DNA was extracted from the iPSC TH-Rep1-SNP-BAG3 clone and from one non-edited clone according to the supplier’s protocol (DNeasy Blood and Tissue, QIAGEN). The BAG3 SNP-containing region of interest was then amplified by PCR using the primers SNP_BAG3_gDNA_F and SNP_BAG3_gDNA_R.

The amplified PCR fragment was purified on agarose gel (MinElution gel extraction kit QIAGEN) and sent for sequencing by Eurofins.

#### Stemness markers analysis on iPSC TH-Rep1-SNP-BAG3 clone by RTqPCR

Relative gene expression analysis of pluripotency markers (relative to the housekeeping gene ACTB) was performed by RTqPCR on iPSC TH-Rep1-SNP-BAG3 clone (SNP-BAG3) and compared to iPSC TH-Rep1 (WT). The following sets of primers were used: NANOG_F and NANOG_R; OCT3/4_F and OCT3/4_R; DNMT3B_F and DNMT3B_R; ACTB_F and ACTB_R.

#### Stemness markers analysis on iPSC TH-Rep1-SNP-BAG3 clone by immunocytochemistry

Expression of stemness markers was analyzed by immunocytochemistry. iPSC TH-Rep1-SNP-BAG3 cells were stained with primary antibodies mouse monoclonal OCT3/4 (C-10) (sc-5279) or mouse monoclonal SOX2 (E-4) (sc-365823) and a secondary antibody Goat anti-Mouse IgG (H+L), Alexa Fluor^™^ 647 (Thermofisher Scientific A-21235), and DAPI. Images were acquired using BC43 Andor microscope, objective X40 (scale bar: 30 um).

#### Stemness and neural progenitor markers analysis during neural induction of iPSC into NPC TH-Rep1-SNP-BAG3 by RTqPCR

Relative gene expression (Log2 Fold change) analysis of stemness and neural progenitor markers was performed by RTqPCR on NPC TH-Rep1-SNP-BAG3 and normalized to iPSC TH-Rep1-SNP-BAG3. The following sets of primers were used: Stemness markers: NANOG_F and NANOG_R; OCT3/4_F and OCT3/4_R. Neural progenitor markers : NESTIN_F and NESTIN_R; PAX6_F and PAX6_R; SOX1_F and SOX1_R. Housekeeping gene: ACTB_F and ACTB_R.

Brightfield images of NPC TH-Rep1-SNP-BAG3 (passage 5) and differentiated TH-Rep1-SNP-BAG3 neurons at day 7 showing their typical morphologies.

#### Microscopy images of differentiated mDANs TH-Rep1-SNP-BAG3

Differentiated mDANs (Day 16) were stained with primary antibodies rabbit polyclonal anti-Tyrosine Hydroxylase (TH) Antibody (Sigma Ab152) and chicken polyclonal anti-MAP2 (AbCam ab5392) with secondary antibodies goat anti-rabbit IgG (H+L), Alexa Fluor™ 647 (Thermofisher Scientific A-21245) and goat anti-chicken IgY (H+L), Alexa Fluor™ 488 (Thermofisher Scientific A-11039) and DAPI. Images were acquired using BioTek Cytation C10 Confocal Imaging Reader (Agilent), objective X20 (scale bar : 200 um).

### Flow cytometry

Cell media was removed, and cells were incubated 5 minutes with accutase (Sigma-Aldrich, A6964) at 37°C. Accutase was deactivated by adding twice the volume of DMEM/F-12 and cell dissociation was further carried on by pipeting the cell suspension. Next, the cell suspension was passed through a DMEM/F-12 primed 50µm cell strainer (Sysmex Partec-Celltrics, 04-004-2327) to ensure a homogenous single-cell suspension. Cells were pelleted at 300x*g* for 5 minutes at 20°C and the pellet was then resuspended in phosphate buffer saline (PBS) (Gibco, ThermoFisher Scientific, 14190185). Sytox blue (ThermoFisher, S34857) was added at a dilution of 1:1000 and cells were incubated for 5 minutes at room temperature to stain dead cells. Neurons incubated at 70°C for 5 minutes were used as positive control for Sytox blue. The flow cytometry analysis was performed using a BD FACSMelody.

Differentiated mDANs TH-Rep1-SNP-BAG3 (day 17) expressing the mCherry Tyrosine Hydroxylase reporter were analyzed by flow cytometry. Differentiated 17608/6 (mCherry negative) and TH-Rep1 mDANs (mCherry positive) were quantified. FlowJo version 10 software was used for data processing and generation of plots.

### Total RNA extraction, cDNA synthesis and RT-qPCR

Total RNA was extracted from TH-Rep1 neurons using the Quick-RNA Microprep kit (Zymo Research, R1051) according to the manufacturer’s instructions.

cDNA synthesis was performed using between 250 ng and 500 ng of total RNA, 0.5 mM dNTPs (ThermoFisher Scientific, R0181), 2.5 µM oligo dT(18)-primer (Eurogentec, Belgium), 1 U/µL Ribolock RNase inhibitor (ThermoFisher Scientific, EO0381) and 5 U/µL RevertAid Reverse transcriptase (ThermoFisher Scientific, EP0442) for 1h at 42°C. The PCR reaction was stopped by incubating samples at 70°C for 10 minutes.

RT-qPCR was accomplished in an Applied Biosystems 7500 Fast Real-Time PCR System using Thermo Scientific Absolute Blue qPCR SYBR Green Low ROX Mix (ThermoFisher Scientific, AB4322B). Each reaction consisted of 5 µL of cDNA, 5 µL of primer pairs (2 µM) and 10 µL of the Absolute Blue qPCR mix. PCR reactions were achieved using the following conditions: 95°C for 15 minutes to activate the DNA polymerase followed by 40 cycles of 95°C for 15 seconds, 55°C for 15 seconds and 72°C for 30 seconds. Gene expression levels were calculated using the 2^−(ΔΔCt)^ method where ΔΔCt is equal to (ΔCt_(target gene)_ - Ct_(housekeeping gene)_)_tested condition_ - (ΔCt_(target gene)_ - Ct_(housekeeping gene)_) _control condition_ and *ACTB* was used as the housekeeping gene. Sh*SCRAMBLE* was used as control condition.

### ATAC-seq experiments

The ATAC-seq protocol was performed with 60 000 iPSCs or NPCs of TH-Rep1-SNP-BAG3 cell line according to (22). Cells were detached with Accutase. For library quality control, the Agilent High Sensitivity DNA kit was used in a 2100 Bioanalyzer instrument. Libraries were sequenced on a NextSeq2000 machine using paired-end 50 bp read length.

### ATAC-seq data analysis

Raw fastq files were assessed for quality using FastQC/fastq_screen. A summary of sample quality control was obtained using MultiQC. Our snakemake template (https://gitlab.com/uniluxembourg/fstm/dlsm/bioinfo/snakemake-atac_seq) (22), where the software set is bundled in a Docker image published here: https://hub.docker.com/r/ginolhac/snake-atac-seq tag 0.3. Trimming of sequencing adapters was done using AdapterRemoval (minimum length 35 bp and adapter sequence: CTGTCTCTTATACACATCT) and mapping to the reference genome with Bowtie2. Quality reads (≥Q30) were filtered using SAMtools. Genome version was GRCh38 Ensembl release 102.

### Allelic imbalance in chromatin accessibility

Allelic imbalance in chromatin accessibility was performed using the R package AllelicImbalance (25) version 1.36.0 in R 4.2.3 and Rstudio (v2023.12.1.402). Briefly, heterozygous sites in the ATAC-seq BAM files of smNPCs, mDAN D15, mDAN D30, mDAN D50 and astrocytes D65 derived from TH-Rep1 cell line, and iPSCs and smNPCs from TH-Rep1-SNP-BAG3 cell line were quantified. The allele count was then calculated, and a chi-square test or generalized linear mixed model (random effect for the replicate) using the binomial distribution were computed as indicated to assess statistical significance between effect and the other allele.

### Cell type association with genetic risk of Parkinson’s disease

To evaluate whether common SNPs associated with increased risk of PD in the European population are enriched in sorted cell type-specific expressed genes, we used Multi-marker Analysis of GenoMic Annotation (MAGMA) v1.09. MAGMA is a gene set enrichment analysis method that tests the joint association of all SNPs in a gene with the phenotype while accounting for the LD structure between SNPs (26). In our study, the SNPs and their corresponding p-values were taken from the summary statistics of the PD GWAS from Nalls *et al*. 2019 (17). LD between SNPs was estimated using the publicly available European subset of 1000 Genomes Phase 3 as a reference panel. The MAGMA analysis comprises three steps. The first step is the annotation process, where SNPs are mapped to genes using the NCBI GRCh37 build (annotation release 105). Gene boundaries are defined as the transcribed region of each gene. A 35 kb window upstream and 10 kb window downstream of each gene were added to the gene boundaries. Next, the gene analysis step calculates gene-wise p-values based on SNP p-values. The third step is the competitive gene set analysis implemented as a linear regression model on a gene set data matrix. The gene sets used in this study consist of differentially expressed genes (>1.33 fold over the median expression given as transcripts per million or TPM) in each sorted cell type. The MAGMA gene set analysis provides association results for each gene set and all genes within the gene sets.

### Identification of SNPs that affect TF binding

To identify regulatory SNPs that affect TF binding sites and therefore may change target gene expression, we applied our software SNEEP ((27), GitHub: https://github.com/SchulzLab/SNEEP). The software requires as input a list of SNPs and a set of TF motifs. For each resulting SNP-TF motif pair, SNEEP evaluates whether the binding behaviour of the TF is affected by the analysed SNP.

For each timepoint of differentiation (D15 pos, D30 pos, D50 pos, and smNPC), we analysed the SNPs that overlap with the timepoint-specific accessible regions. For the accessible regions we took the ATAC peaks from our previous work (22). As TF motifs we used 633 non-redundant human motifs from the JASPAR database (version 2020) (28), where we excluded for each timepoint of differentiation the non-expressed TFs (counts <=0.5).

SNEEP incorporates the functionality to consider SNPs only in regions of interest (in our case within the accessible regions) and excluding TFs not or weakly expressed. To do so, we specified the following parameter for each timepoint of differentiation: -p 0.05, -c 0.01 -f <ATAC-peaks-timepoint-specific> - t <RNA-seq-timepoint-specific> -e <MappingEnsemblIDToGeneNameForTFs> -d 0.5. Additionally, the PD GWAS SNPs and the TF motifs are required to run our software.

To find the genes affected by the SNPs identified by SNEEP, the timepoint-specific accessible-regions had to be linked to their target genes. This was done with STARE (v1.0.4) (29) (GitHub: https://github.com/SchulzLab/STARE)). For the candidate enhancers the narrowPeaks ATAC-seq files of each timepoint were used and the signalValue taken as enhancer activity. As gene annotation we took v38 from GENCODE (30). The LowC data was processed with FAN-C (v.0.9.27) (31) and used as contact data for STARE at 5 kb resolution. All other parameters were kept at their default values (window size 5MB, score-cutoff 0.02).

### Primers

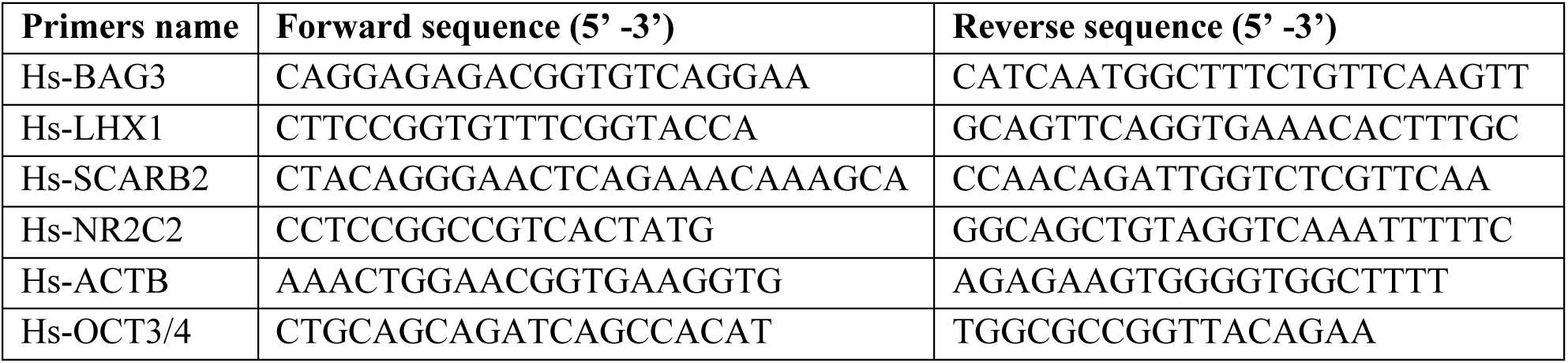
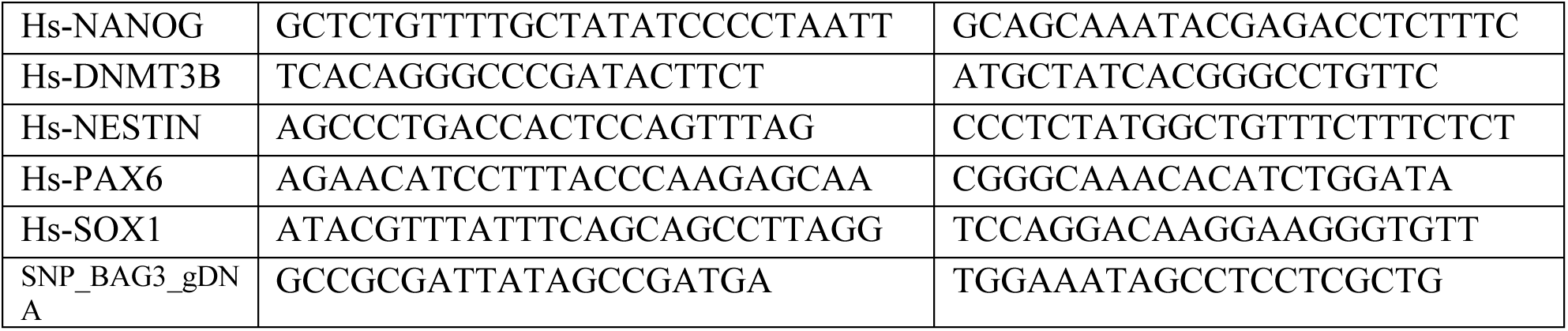

### Prime editing oligonucleotides

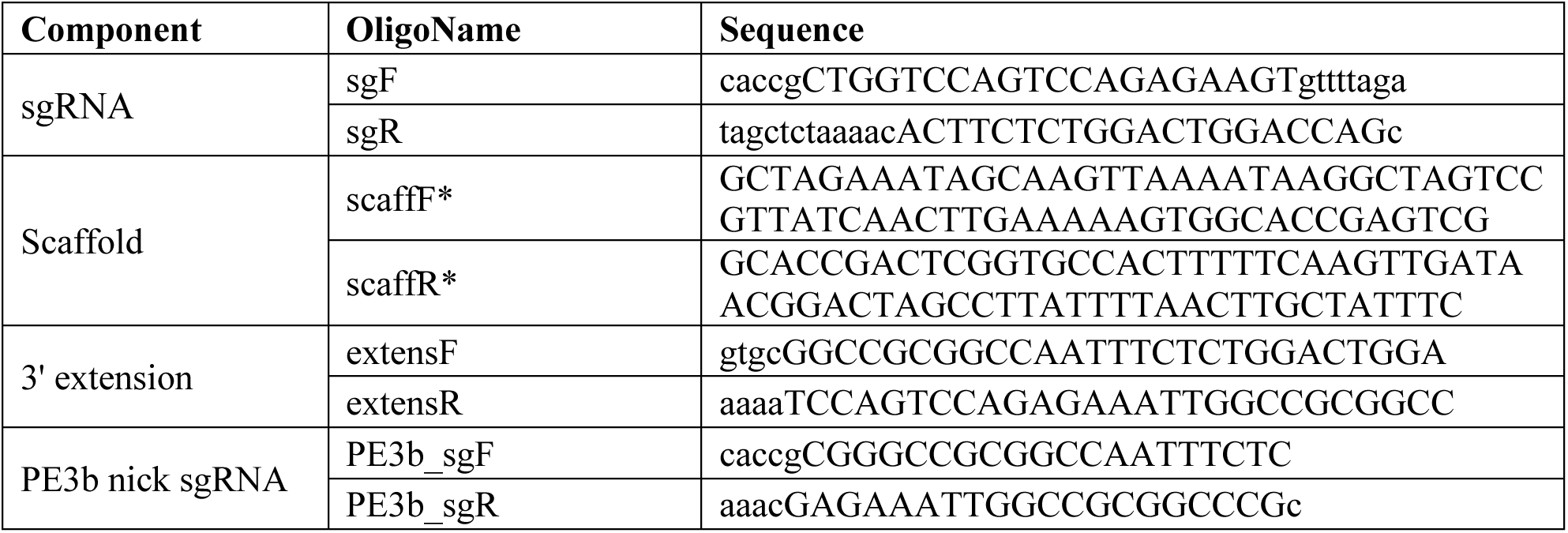

### Bacterial glycerol stocks

Bacterial glycerol stocks were purchased from GeneCopoeia (LabOmics). The different shRNA and luciferase constructs used were:

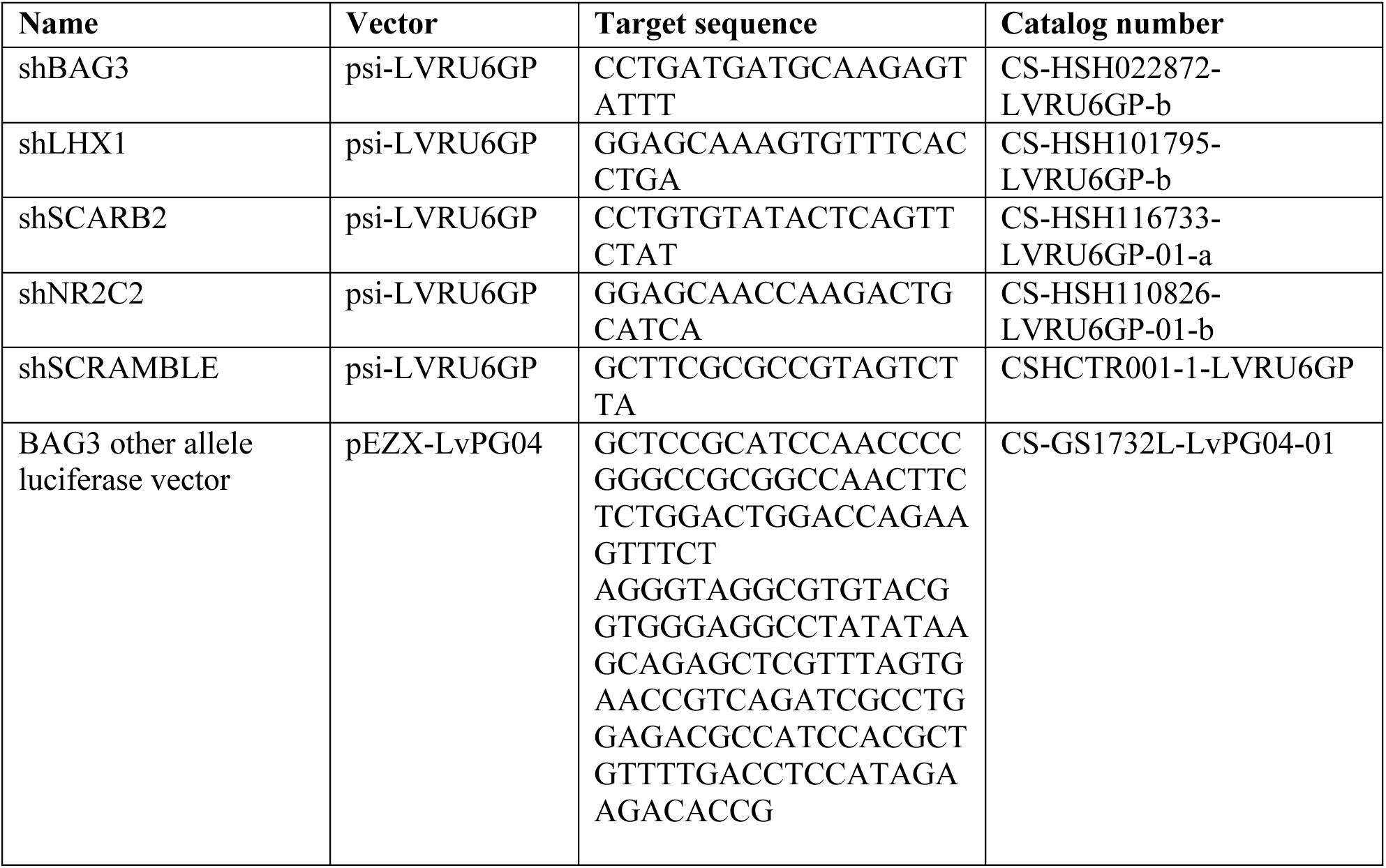
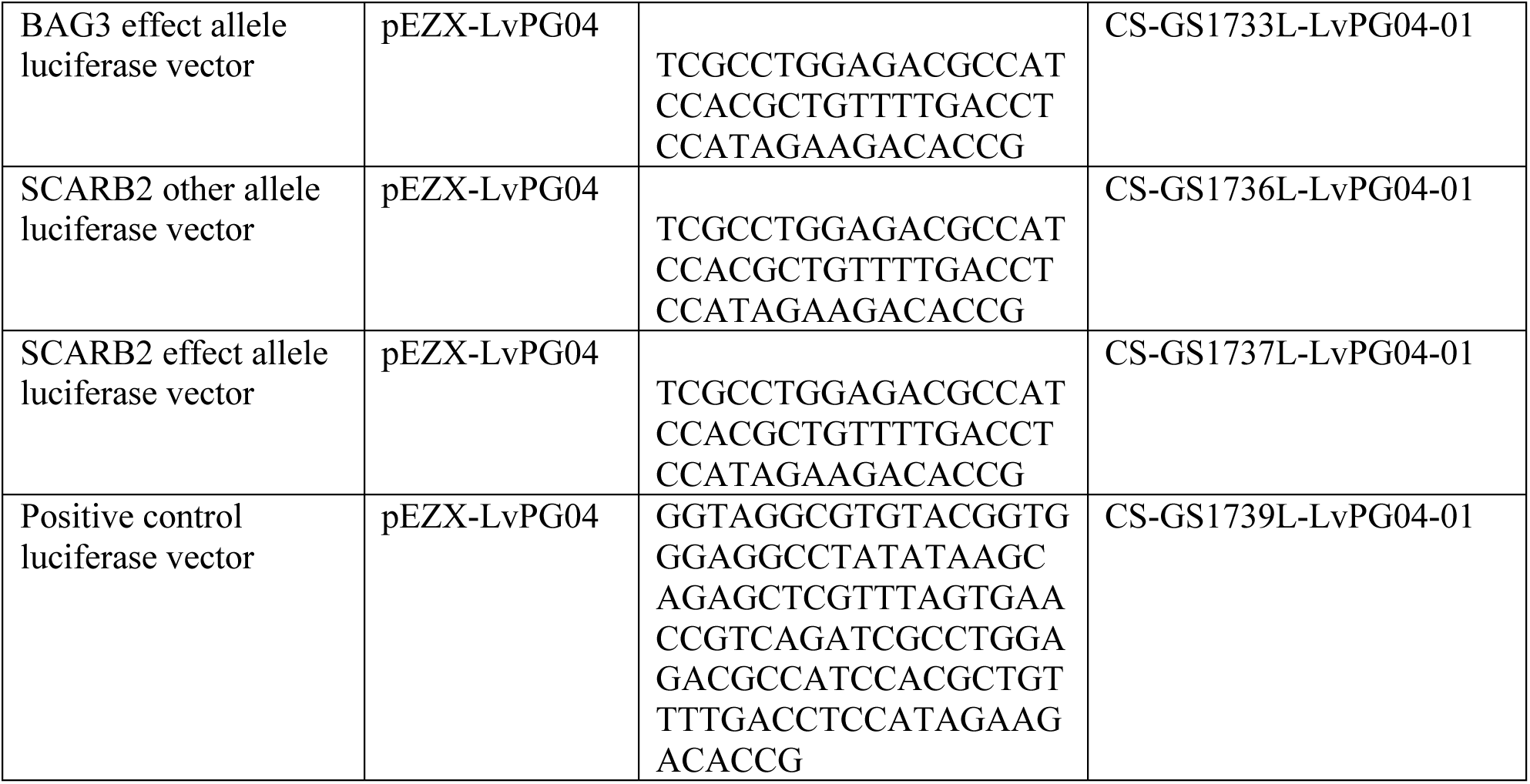

### Low chromosome capture (LowC)

#### Cell Fixation and Lysis Preparation

After cell sorting, 110 000 TH-Rep1 smNPCs or mDANs differentiated for 30 days were transferred into a new 1.5mL tube and were centrifuge for 5 minutes at 300 x *g* at room temperature. Following centrifugation, the supernatant was removed and the pellet was resuspended in 946 µl of Dulbecco’s phosphate buffer saline (DPBS) (Gibco, ThermoFisher Scientific, 14190169). Then, samples were cross-linked with formaldehyde (Sigma-Aldrich, 1.03999) at a final concentration of 1% for 10 minutes at room temperature on a rotating wheel. The cross-linked reactions were then quenched with ice-cold glycine (Carl Roth, 3908.3) at a final concentration of 0.2 M for 5 minutes at room temperature on a rotating wheel. Immediately after, samples were centrifuged at 300 x *g* for 5 minutes at 4°C. Following centrifugation, the supernatant was removed and the pellets were resuspended in ice-cold DPBS (Gibco, ThermoFisher Scientific, 14190169) and centrifuge again at 300 x *g* for 5 minutes at 4°C. The previous washing step was again performed and the cells were lysed in 250 µl of ice-cold *in situ* Hi-C lysis buffer [10 mM Tris-HCl (VWR, A1086.1000) pH 8.0, 10 mM NaCl (Carl Roth, 3957.1), 0.2% IGEPAL CA-630 (Sigma Aldrich, I8896), autoclaved distilled water] containing mini protease inhibitor cocktail (PI) (Roche, 4693124001) and incubated on ice for 15 minutes.

#### Nuclei Isolation and SDS Permeabilization

The cell lysates were then centrifuged at 1000 x *g* for 5 minutes at 4°C. Following centrifugation, the supernatants were discarded and cells were resuspended again in 125 µl of ice-cold *in situ* Hi-C lysis buffer. The samples were centrifuged at 13000 x *g* for 5 minutes at 4°C followed by the removal of the supernatant and resuspension in 250 µl of ice-cold 10X NEB2 buffer [500 mM NaCl (Carl Roth, 3957.1), 100 mM Tris-HCl (VWR, A1086.1000) pH 8.0, 100 mM MgCl_2_ (Sigma-Aldrich, M8266), 10 mM dithiothreitol (DTT) (Sigma-Aldrich, D9779), autoclaved distilled water]. The cell lysates were again centrifuged at 13000 x *g* for 5 minutes at 4°C and the supernatant were discarded. To further lyse the nuclei, the cell pellets were resuspended in 25 µl of 0.4% sodium dodecyl sulfate (SDS) (Carl Roth, 2326.1) and immediately incubated at 65°C for 10 minutes. SDS was quenched by adding 12.5 µl of 10% Triton X-100 (Sigma-Aldrich, X100) and 50 µl of nuclease-free water and incubated at 37°C for 45 minutes in a thermomixer at 650 rpm.

#### Restriction Digest, End Filling, and Proximity Ligation

DNA digestion was performed by adding 20 µl of 10X NEB2.1 buffer with 15 µl of MboI restriction enzyme (NEB R0147L, New England Biolabs) and incubated at 37°C for 45 minutes in a thermomixer at 450 rpm. After 45 minutes, 5 µl of MboI restriction enzyme (NEB R0147L, New England Biolabs) was added again and incubated at 37°C for 45 minutes in a thermomixer at 450 rpm to increase digestion efficiency. Inactivation of the MboI restriction was performed by incubating the samples at 62°C for 20 minutes. To fill the restriction fragment overhangs, 18.75 µl of 0.4mM biotin-14-dCTP (19518018, ThermoFisher), 0.75 µl of 10 mM dATP (U120D, Promega), 0.75 µl of 10 mM dGTP (U121D, Promega), 0.75 µl of 10 mM dTTP (U123D, Promega) and 8 µl of 5U/µl DNA polymerase I Klenow (NEB M0210L, New England Biolabs) was added and the samples were incubated at 37°C for 90 minutes in a thermomixer at 400 rpm. The step of DNA ligation was set up by adding 657 µl of nuclease-free water, 120 µl of 10X T4 DNA Ligase buffer (EL0011, ThermoFisher), 100 µl of 10% Triton X-100 (Sigma-Aldrich, X100), 12 µl of 20 mg/ml bovine serum albumin (BSA) (Carl Roth, 8076.4) and 5 µl of 5 Weiss U/µl T4 DNA ligase (EL0011, ThermoFisher) to the samples that were incubated at room temperature for 4 hours on a rotating wheel. After this incubation time, the samples were centrifuged at 2500 x *g* for 5 minutes at room temperature.

#### Reverse Cross-linking and DNA Purification

For protein digestion and reverse cross-linking, the supernatant was discarded and the pellet was resuspended in 500 µl of extraction buffer (50 mM Tris-HCl (VWR, A1086.1000) pH 8.0, 50 mM NaCl (Carl Roth, 3957.1), 1 mM EDTA (Carl Roth, CN06.3), 1% SDS (Carl Roth, 2326.1), autoclaved distilled water) and 20 µl of Proteinase K (EO0491, ThermoFisher) and incubated at 55°C for 30 minutes in a thermomixer at 1000 rpm followed by the addition of 130 µl of 5M NaCl (Carl Roth, 3957.1). The samples were then incubated overnight at 65°C in a thermomixer at 1000 rpm. DNA extraction was performed by adding 630 µl of phenol-chloroform-isoamyl alcohol mixture (25:24:1) (Sigma-Aldrich, 77617) and the samples were vortexed at max speed for one minute followed by centrifugation at 17000 x *g* for 5 minutes at 4°C. Then, the supernatant was transferred to a new tube and precipitation of DNA was carried on by adding 63 µl of 3M sodium acetate (C_2_H_3_NaO_2_) (Sigma-Aldrich, 71183), 2 µl of GlycoBlue (Invitrogen, ThermoFisher, AM9515) and 1008 µl of absolute ethanol (Carl Roth, 9065.2) and incubated at −80°C for 15 minutes. Then the samples were centrifuged at 17000 x *g* for 30 minutes at 4°C and the supernatant was removed. The DNA pellet was washed with 800 µl of 70% ethanol (Carl Roth, 9065.2) pre-cooled at −20°C followed by centrifuging the samples at 21130 x *g* for 5 minutes at 4°C. The supernatant was again removed the pellet was air dried for 5 minutes. Dissolution of DNA pellets was performed by adding 30 µl of 10 mM Tris pH 7.5 (pre-warmed at 37°C) (VWR, A1086.1000). RNA contamination was removed by adding 1 µl of RNAse A (EN0531, ThermoFisher) and incubated at 37°C for 15 minutes.

#### Biotin Removal and Chromatin Fragmentation

Next, the removal of biotin from unligated fragments was performed using 5 µg of DNA together with 10 µl of 10X NEBuffer r.2.1 (NEB M0203S, New England Biolabs), 2.5 µl of 1 mM dATP (U120D, Promega), 2.5 µl of 1 mM dGTP (U121D, Promega), 2.5 µl of 1 mM dTTP (U123D, Promega), 2.5 µl of 1 mM dCTP (U122D, Promega), 0.5 µl of BSA 20 mg/ml (Carl Roth, 8076.4), 5 µl of 3U/µl T4 DNA polymerase (NEB M0203S, New England Biolabs) and 45.5 µl of nuclease-free water followed by an incubation at room temperature for 4 hours. Biotinylated DNA samples were brought to 120 µl with nuclease-free water in a low-binding 1.5 ml tubes and were sheared with a sonicator (Bioruptor Pico, Diagenode) during 6 cycles (30 seconds ON and 90 seconds OFF).

#### Streptavidin Pulldown of Biotinylated DNA

For the pulldown of biotinylated fragments, 30 µl of Dynabeads MyOne Streptavidin C1 (Invitrogen, ThermoFisher, 65001) beads per sample were washed with 400 µl of of 1X Bind & Wash buffer [5 mM Tris-HCl pH 7.5 (VWR, A1086.1000), 0.5 mM EDTA (Carl Roth, CN06.3), 1 M NaCl (Carl Roth, 3957.1), 0.05% Tween 20 (Sigma-Aldrich, P1379), autoclaved distilled water] added with 0.1% Triton X-100 (Sigma-Aldrich, X100). After separation on a magnet using a DynaMag^TM^-2 magnetic stand (Life Technologies, 12321D), the supernatant from the beads was discarded and the beads were resuspended in 30 µl per sample of 2X Bind & Wash buffer [10 mM Tris-HCl pH 7.5 (VWR, A1086.1000), 1 mM EDTA (Carl Roth, CN06.3), 2 M NaCl (Carl Roth, 3957.1), autoclaved distilled water]. Then, 30 µl of washed beads were added to the sheared DNA and the samples were incubated at room temperature for 15 minutes on a rotating wheel. The beads were then captured using a DynaMag^TM^-2 magnetic stand (Life Technologies, 12321D) and the supernatant was removed. The beads were then washed twice in 400 µl of 1X Bind & Wash buffer added with 0.1% Triton X-100 (Sigma-Aldrich, X100) and incubated at 55°C for 2 minutes in a thermomixer at 650 rpm. The beads were again captured and the supernatant was discarded. Finally, the beads were washed once with 400 µl of 1X NEB2 buffer [50 mM NaCl (Carl Roth, 3957.1), 10 mM Tris-HCl (VWR, A1086.1000) pH 8.0, 10 mM MgCl_2_ (Sigma-Aldrich, M8266), 1 mM DTT (Sigma-Aldrich, D9779), autoclaved distilled water]. Next, the beads were captured on the DynaMag^TM^-2 magnetic stand (Life Technologies, 12321D) and the supernatant was discarded.

#### End Repair, Adapter Ligation, and Indexing

For end repair, the beads were resuspended in 55.5 µl of 10 mM Tris pH 8.0 (VWR, A1086.1000) and 3 µl of End Prep enzyme mix with 6.5 µl of End repair reaction buffer were added (NEB E7645S, New England Biolabs). The reaction was carried on in a Analytik Jena Biometra thermocycler with the heated lid ON and the following program: 30 minutes at 20°C, 30 minutes at 65°C and hold at 4°C. Addition of indexes was performed by adding 15 µl of Blunt/TA Ligase master mix (NEB E6440S), 2.5 µl of NEBNext Adaptor (NEB E6440S) and 1.0 µl of ligation enhancer (NEB E6440S) and was incubated at 20°C for 15 minutes in a Analytik Jena Biometra thermocycler. Next, 3 µl of USER enzyme (NEB E6440S) was added to the reaction and was incubated at 37°C for 15 minutes in a Analytik Jena Biometra thermocycler. The beads were next washed with 200 µl of 1X Bind & Wash buffer added with 0.1% Triton X-100 (Sigma-Aldrich, X100) and transferred to a new tube. The beads were captured by a DynaMag™-2 magnetic stand (Life Technologies, 12321D) and the supernatant was removed. The wash was repeated and the beads were then wash once with 200 µl of 10 mM Tris pH 8.0 (VWR, A1086.1000) and transferred to a new tube. Finally, beads were captured in a magnet and the supernatant was removed followed by the resuspension of the beads in 50 µl of 10 mM Tris pH 8.0 (VWR, A1086.1000).

#### Library Amplification, Size Selection, and Quality Control

Finally, 4 amplification runs were run in parallel to maximise library diversity by adding to 10 µl sample, 25 µl of 2X NEBNext Q5 Hot Start HiFi PCR Master mix (NEB 7645S), 5 µl Unique Dual Index Pairs (NEB6440S) and 10 µl nuclease-free water. PCR enrichment was performed using the following program in a Analytik Jena Biometra thermocycler: 98°C for 1 minute, 14 cycles at 98°C for 10 seconds followed by 65°C for 75 seconds (ramping 1.5°C per second), 65°C for 5 minutes and hold at 4°C. Following PCR enrichment, the four reactions were combined. Size selection was performed with SPRI select beads (Beckman coulter, B23318) according to the manufacturer instructions. Quality control was assessed with the Agilent High Sensitivity DNA kit (Agilent, 5067-4626).

### LowC data analysis

The tool suite HiCExplorer (v3.7.6) (32) was used for mapping, generating pairs files and binned LowC matrices as h5 files, one per replicate. Those matrices were further merged per condition and converted to cool files for downstream processing using FAN-C (v0.9.23b) (31) to investigate PC1 and TADs. Calculation of intra- or inter-compartment strengths were done from the cool files at resolution 500 kb using pentad (https://github.com/magnitov/pentad). Plotting LowC data was perfomed with the R package plotgardener (v1.4.2) (33) in R statistical software (v4.2.3) (34).

## Results

### PD-associated variants are enriched in vicinity of genes expressed in iPSC-derived mDANs

To investigate whether non-coding PD SNPs identified in GWAS can contribute to PD risk by acting as regulatory SNPs in a cell type-specific manner, we took advantage of our recent multiomic data of iPSC-derived cultures of astrocytes and dopaminergic neurons (mDANs) (Figure 1A, (22)). The data consist of paired transcriptomic and chromatin accessibility data from neural progenitor cells (smNPCs), astrocytes and different time points of differentiation of purified reporter-positive dopaminergic neurons (mDANs) and reporter negative non-mDANs. The presence of an integrated mCherry reporter at TH locus enabled the purification of mDANs from mixed neuronal cultures by FACS (21). Inspection of established markers genes of mDANs, including known transcriptional regulators and enzymes and transporters involved in dopamine metabolism, confirmed their selective induction and expression in mCherry positive mDAN population (Supplementary Figure S1). Recent single cell transcriptomic analysis of all cell types of human substantia nigra have demonstrated a specific enrichment of PD SNPs in proximity of genes expressed in mDANs *in vivo* (35,36). Therefore, using our transcriptomic data, we selected genes with cell type-selective expression for each cell type and differentiation state and used MAGMA (26) to determine whether the PD SNPs from GWAS meta-analysis (17) are enriched in vicinity of the genes in these sets also *in vitro* (Figure 1B). Interestingly, PD SNPs were significantly enriched in proximity of genes showing selective expression in mDANs, suggesting they might also play an important role in *cis*-regulatory variation in these cells.

**Figure 1:**
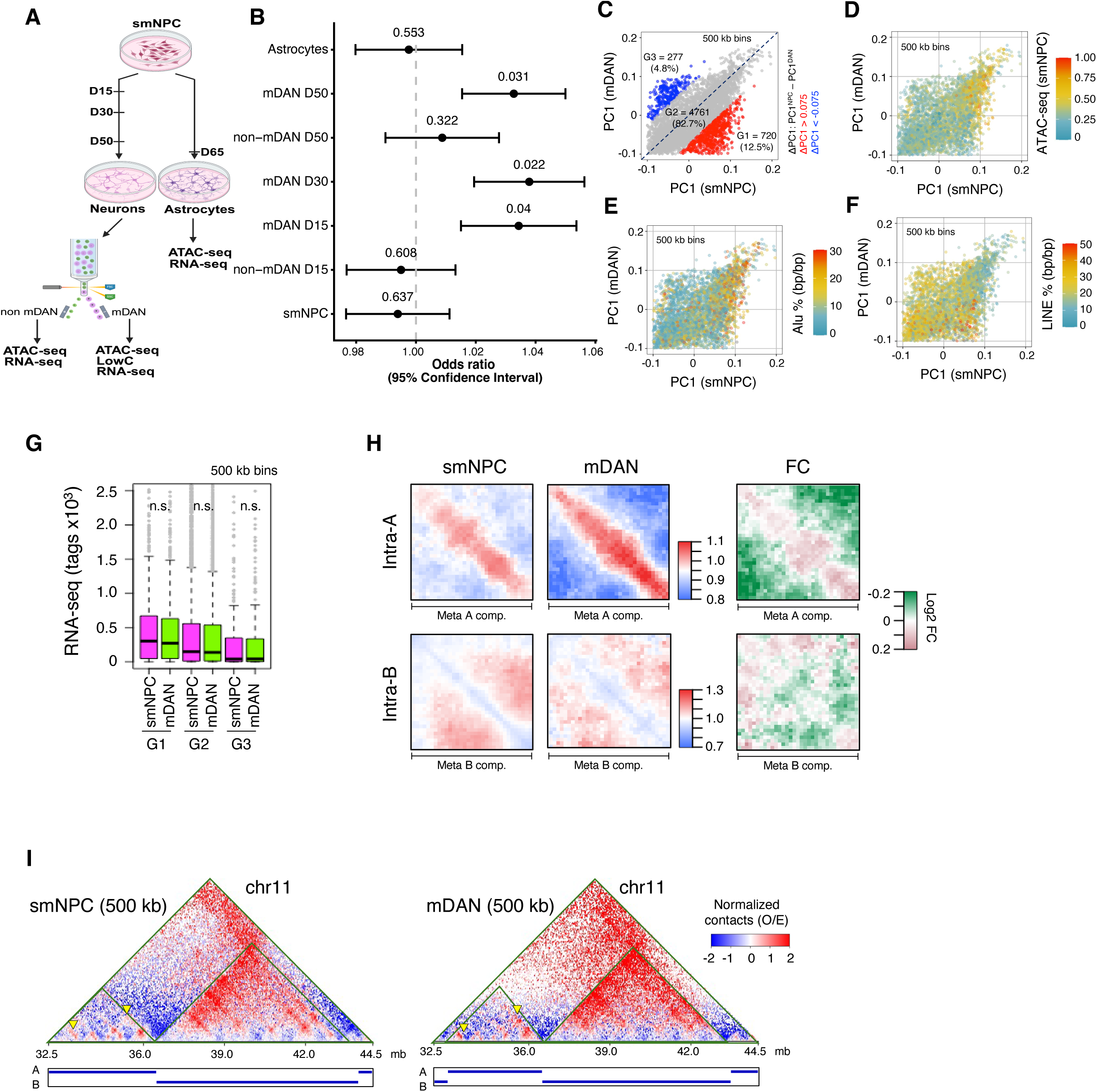
3D genome organization in smNPCs and mDANs. **A**) Schematic representation of the experimental set-up. TH-Rep1 cell line was differentiated into neurons and astrocytes. Neurons were submitted to fluorescense activated cell sorting (FACS) to isolate dopaminergic neurons (mDAN) from non-dopaminergic neurons (non-mDAN). Total RNA and chromatin were collected at the indicated time points and subjected to RNA-seq, ATAC-seq, and LowC analysis. **B)** Forest plot of the odds ratio of PD rSNPs in vicinity of genes expressed in mDAN, non-mDAN and astrocytes. **C)** Dot plot showing PC1 values for smNPCs and mDANs, calculated at 500 kb resolution. Red dots denote 500 kb bins that showed a decrease in PC1 values in mDANs (PC1smNPC – PC1mDAN > 0.075, group G1), while blue dots denote 500 kb bins that showed a reciprocal increase in PC1 values in mDANs (PC1smNPC – PC1mDAN < −0.075, group G3). The rest are shown as gray dots (group G2). **D)** Dot plot showing normalized ATAC-seq read depth in 500 kb genomic bins. Genomic bins showing high chromatin accessibility (high ATAC-seq read depth) are denoted in red, while those showing low chromatin accessibility (low ATAC-seq read depth) are denoted in blue. **E)** Dot plot showing percentage (bp/bp) of Alu elements in 500 kb genomic bins, in smNPCs and mDANs. Alu-rich or -poor regions are denoted in red or blue, respectively. **F)** Dot plot showing percentage (bp/bp) of LINE elements in 500 kb genomic bins, in smNPCs and mDANs. LINE -rich or -poor regions are denoted in red or blue, respectively. **G)** RNA expression levels shown for G1 to G3 groups of bins, for smNPCs and mDANs. There were no significant statistical differences observed by Student’s t-test. **H)** Metaplots showing intra-A (top) and intra-B (bottom) interactions. The color code denotes interaction levels between neighboring regions (red = strong, blue = weak). A plot based on the fold change (FC) of values between smNPCs and mDANs is shown at the right for intra-A or intra-B interactions. Stronger interactions in smNPCs are shown in magenta, while weaker interactions are shown in green. **I)** A representative region (chr11: 32.5 mb – 44.5 mb) showing a gain in intra-A interactions, denoted by inverted yellow triangles(red= increased contacts, blue = decreased contacts).

### Higher order genome organization in smNPCs and mDANs

Rewiring of the higher-order genome organization is associated with neural development and differentiation (37–39). Recent genome-wide chromatin conformation capture analyses such as Hi-C have revealed that the higher-order genome in mammalian cells is spatially segregated into megabase sized euchromatic or heterochromatin domains known as A or B compartments (40). To annotate higher-order genome organization in smNPCs and mDANs, and to allow more accurate association of the identified enhancers to their target genes in smNPCs and purified mDANs, we carried out LowC experiments to define the long-distance chromatin interactions and topologically associated domains (TADs) present in these cells.

We calculated PC1 (principle component 1) of the eigenvector of the LowC matrices and called A (PC > 0) or B (PC1 < 0) compartments with a resolution of 500 kb (Figure 1C). Previous studies have reported that A or B compartments are enriched in SINE/Alu (short interspersed nuclear elements in mice, Alu in humans) or LINE retrotransposons, respectively (41). Indeed, our analysis confirmed that in smNPCs and mDANs, A compartments exhibited higher mean accessibility as measured by ATAC-seq, and were enriched for Alus, while B compartments exhibited enrichment for LINEs (Figure 1D-F). Visual inspection of a representative chromosome (chr 11) further corrobarated these findings by showing that Alu or LINE repeats were enriched in A or B compartments, respectively (Supplementary Figure S2).

We next tested whether differentiation from smNPCs to mDANs was associated with structural changes in the higher order genome organization, by plotting PC1 values in 500 kb genomic bins for smNPCs and mDANs (Figure 1C). This analysis showed that 12.5% of all 500 kb bins showed a decrease in mDANs (group G1), while 4.5% showed a reciprocal increase (group G3). The rest (82.7%, group G2) remained unchanged (Figure 1C). A decrease in PC1 values indicates a shift to a more heterochromatin-like state. A considerable fraction of the 500 kb genomic regions, therefore, may exhibit increased heterochromatin-like properties upon differentiation into mDANs. A recent study showed that 3D organization of the genome is different between neuronal and non-neuronal cells (39). We mined the Hi-C data from this report, and calculated A/B compartments in non-neuronal (NeuN negative: NeuN^neg^) or neuronal (NeuN positive: NeuN^pos^) post-mortem brain cells (Supplementary Figure S3). However, we found A/B compartments to remain largely stable between NeuN^neg^ and NeuN^pos^ cells, with somewhat lower number of changes to those observed in mDAN differentiation (Supplementary Figure S3A). Taken together, our results indicate that the differentiation of smNPCs to mDANs may be accompanied by a significant reorganization of the 3D genome, as manifested by the decrease of PC1 values in 12.5% of all 500 kb genomic bins.

We wondered whether the changes in the 3D genome organization were also associated with changes in transcription during differentiation into mDANs (Figure 1G). We compared the transcription levels in the 500 kb genomic bins between smNPCs and mDANs. The G1 group of bins that possess higher PC1 values and are therefore more euchromatic, generally exhibited higher transcription levels. In contrast, the G3 group of bins that possess lower PC1 values and are therefore more heterochromatic, also exhibited lower transcription levels. However, direct comparison between smNPCs and mDANs within each group of regions (G1, G2 or G3) showed no clear changes in transcription levels (Figure 1G). To examine whether there were any changes in chromatin accessibility between smNPCs and mDANs, we used our ATAC-seq data to compare the groups of regions (Supplementary Figure S2B). Consistent with the expression levels (Figure 1G), euchromatin regions with higher PC1 values exhibited more chromatin accessibility, while heterochromatic regions with lower PC1 values exhibited less chromatin accessibility (Supplementary Figure S2B). Again, there were no significant differences between smNPCs and mDANs. Based on these observations, we concluded that the changes in higher order chromatin organization in mDANs, may not necessarily accompany changes in transcription or chromatin accessibility at genome-wide scale when observed at the resolution of 500 kb bins, consistent with smNPCs being already committed to neuronal lineage.

### Chromatin contacts within and between A/B compartments change upon mDAN differentiation

To further elucidate the changes associated with A or B compartments during mDAN differentiation, we took advantage of a published computational tool (42), and calculated chromatin interactions within A (intra-A) or B (intra-B) compartments (Figure 1H). A compartments exhibit permissive chromatin states, and consistent with this idea, interactions along the diagonal are prominently observed in intra-A compartment plots. Curiously, we noted that interactions along the diagonal in A compartments were markedly increased in mDANs (Figure 1H, top panel, also note the fold change profile at the right). Contacts within the B compartments (intra-B), however, remained largely unchanged (Figure 1H, bottom panel). Visual inspection in a representative locus in chr11 (32.5 mb to 44.5 mb) showed an increase of chromatin contacts within an A compartment (indicated with inverted yellow arrowheads) (Figure 1I). We again compared these results with previously published Hi-C data for neuronal and non-neuronal cells (39). We found that neuronal cells showed increased chromatin interactions along the diagonal in A compartments (i.e. close-range contacts) (Supplementary Figure S3B), in a manner that was reminiscent of our findings in mDANs (Figure 1H). In B compartments however, neuronal cells showed a reciprocal gain of interactions away from the diagonal (Supplementary Figure S3B). Such gains of intra-B interactions were not observed in our data during differentiation of smNPCs to mDANs (Figure 1H).

Homotypic interactions between similar A (A-A) or B (B-B) compartments, likely driven by phase-separation mediated molecular mechanisms, are characteristic features of the higher-order genome. We, therefore, wondered whether A-A or B-B contacts were altered during mDAN differentiation. We plotted A-A contacts, and observed a moderate increase in mDANs compared to smNPCs (Supplementary Figure S2C, top panel). B-B contacts, in contrast, were markedly decreased in mDANs (Supplementary Figure S2C, middle panel). Consistent with this observation, visual inspection in chromatin 11 also revealed that B-B interactions were indeed decreased at multiple genomic loci (Supplementary Figure S2D). Heterotypic interactions between euchromatic A and heterochromatic B compartments are generally suppressed in the 3D genome. Consistent with this idea, we also observed low A-B contacts in smNPCs. Interestingly, such A-B contacts were even more strongly repressed in mDANs (Supplementary Figure S2C, bottom panel). Our analyses, therefore, demonstrates that large scale reorganization of chromatin contacts occur during neuronal differentiation, characterized by higher intra-A and A-A interactions, but lower B-B or A-B interactions in mDANs.

### Number of TADs decreases but TAD length and insulation increase in iPSC-derived mDANs

Reorganization of the chromatin in megabase sized A/B compartments during mDAN differentiation could be linked with finer changes in the sub-compartment level genome organization. Topologically associating domains (TADs) mediate sub-compartment level genomic structures (40). We, therefore, calculated TADs in smNPCs and mDANs using our LowC data (Figure 2). Remarkably, this analysis revealed a dramatic (more than 5-fold) reduction in the number of TADs in mDANs across all three biological replicates. Indeed, while we detected 2142 TADs in smNPCs, this number was reduced to 413 in mDANs (Figure 2A). This decrease was more prominent in the euchromatin, as the number of TADs located in A compartments were reduced by nearly 8-fold from smNPCs (1191) to mDANs (155).

**Figure 2:**
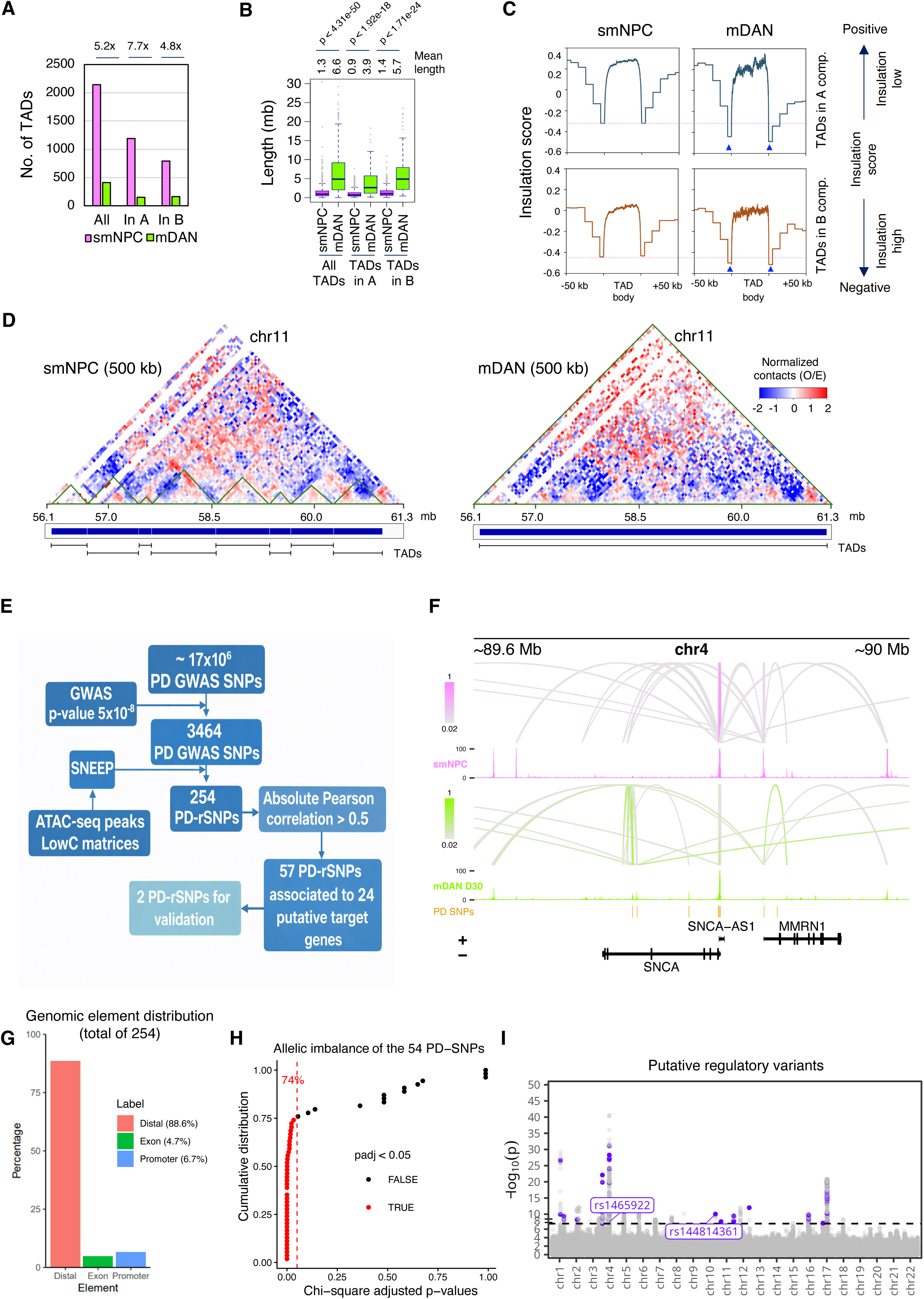
Analysis of chromatin interactions for identification of PD-associated regulatory SNPs (rSNPs). **A)** Number of TADs in smNPCs (pink) or mDANs (green), in total, or in A compartments, or in B compartments, shown as bar plots. The fold change decrease in TAD numbers for each group is shown at the top. **B)** TAD length in smNPCs (pink) or mDANs (green), for all TADs, or TADs in A compartments, or TADs in B compartments, shown as box plots. Median TAD length, and statistical significance (Student’s t-test) of TAD length changes between smNPCs and mDANs, are shown at the top. **C)** Insulation score for TADs is shown in smNPCs (left) and mDANs (right), for TADs in A (top) or B (bottom) compartments. The pink dotted line denotes the insulation level at TAD borders in smNPCs. Note the increased insulation (reduced score) at TAD borders in mDANs (highlighted with blue triangles). **D)** A representative genomic region (chr11: 56.1 mb to 61.3 mb) shows eight separate TADs in smNPCs merging into one large TAD in mDANs. **E)** Schematic representation of the prediction and selection of the studied rSNPs. **F)** Illustration of the predicted regulatory interactions at the *SNCA* locus derived from LowC, ATAC-seq and RNA-seq data using the gABC model. Interactions in smNPCs and day 30 mDANs are shown as lines. The line color indicates the score of the interaction, with higher score indicating higher contribution of the enhancer to the overall regulatory input of the gene. Corresponding ATAC-seq signals are shown below the lines and the location of the predicted PD SNPs are indicated in orange. **G)** Bar plot showing the genomic localization of the 254 putative regulatory PD SNPs, as determined by annotation with the ChIPpeakAnno package (79). Percentages indicate the proportion of SNPs located in distal, exonic, and promoter regions (±1 kb). **H)** Statistical significance of allelic imbalance induced by each of the 54 PD rSNPs present in the used cell line is shown. Each PD SNP had its imbalance assessed by a chi-square test and the smallest p-value across the time points was kept. FDR were computed using the Benjamini-Hochberg method. **I)** Manhattan plot of the PD GWAS variants and their genome-wide association p-values (17). Purple dots represent the 254 PD-rSNPs.

Even in the heterochromatic B compartments, however, we observed that the number of TADs decreased by nearly 5-fold from smNPCs (795) to mDANs (167) (Figure 2A). We wondered whether this reduction in the number of TADs was due to merging of smaller TADs into larger ones in mDANs. To this end, we calculated the mean length of all TADs, or TADs in the A compartment, or TADs in the B compartment (Figure 2B). Indeed, we observed that the length of TADs was increased by more than 5-fold in mDANs (mean length 6.6 mb), compared to smNPCs (mean length 1.3 mb) (Figure 2B). Visual inspection of a representative locus (chr11: 56.1 mb – 61.3 mb), also supported this observation, by showing that eight smaller TADs present in smNPCs were merged into a single large TAD in mDANs (Figure 2D). Of note, by comparing 3D genome organization in neural cells and glial cells, a previous study found that local chromatin structures become weaker in the neuronal cells, likely due to widespread decrease in CTCF binding in the genome (38). Paradoxically, the same study also observed that such weak structures often merge together to build larger structures that are flanked by highly insulated chromatin borders strongly bound by CTCF. Reorganization of chromatin structures observed in our study, characterized by merging of smaller TADs into larger ones, are therefore similar to these previous findings (38).

To elucidate chromatin insulation especially at TAD borders, we plotted insulation score over each TAD (TAD body) and 50 kb flanking regions (Figure 2C). As expected, chromatin insulation was high at the TAD border (negative insulation score represents increased chromatin insulation), but was low at the TAD body. TADs in mDANs, located in either A or B compartments, showed slightly increased chromatin insulation at the TAD border compared to smNPCs (Figure 2C). We compared our results with Hi-C analyses performed in non-neuronal and neuronal cells in a previously published paper (39) (Supplementary Figure S3C). Unlike for mDANs, the NeuN positive cells exhibited a weak bias to smaller TAD sizes (1 mb vs 0.8 mb, for non-neuronal vs neuronal cells), and an increased number of TADs (2922 vs 3598, for non-neuronal vs neuronal cells). Thus, the number of TADs decreased, but the length and insulation of TADs increased specifically in mDANs, likely due to merging of smaller TADs into larger ones.

### Identification of putative PD-associated regulatory variants in human dopaminergic neurons

Next, we employed our recently developed tool SNEEP (27) for prediction of PD SNPs that can alter TF binding specifically in mDANs using the 3464 PD SNPs passing the genome-wide significance threshold for PD association (17). Ultimately, together with our LowC data, this allows for the genome-wide identification of TF-enhancer-gene-interactions altered by PD SNPs. To do this, we analysed chromatin regions accessible in mDANs at different time points of differentiation and performed statistical evaluation of the impact of each co-localized PD SNP on the TF binding score of the identified TFBSs (Figure 2E). Applying a stringent cut-off of p < 0.01 for the TF binding score change, we identified a total of 254 putative regulatory PD SNPs predicted to alter TF binding in smNPCs or mDANs (Figure 2E, Supplementary Table S1). Predicted changes included both TFBS that were disrupted and TFBS that were created by the PD-associated effect allele, with motif disruption occurring more often (Supplementary Table S1).

To associate the putative regulatory PD SNPs (and the respective enhancers) to their target genes, we leveraged our context-specific 3D genome organization data (Figure 1) by integrating it with our RNA-seq and ATAC-seq profiles across mDAN differentiation. We identified the cell type-specific enhancer-target gene interactions by applying our previously published generalized Activity-By-Contact (gABC) model (29). To illustrate this, Figure 2F shows cell type–specific enhancer–target gene interactions at the *α-synuclein* (*SNCA*) locus in mDANs and smNPCs for regulatory elements predicted to be affected by PD-associated SNPs. These data depict an overall increase in predicted interaction strengths and the formation of distal intronic enhancers with mDAN-specific regulatory interactions that are potentially altered by regulatory PD SNPs, consistent with previous reports (43).

An analysis of the relative localization of the putative regulatory PD SNPs showed that while 17 of the variants were located within accessible gene promoters (within 1 kb of TSS) and 12 were located within known exons (possibly disrupting coding sequences), a vast majority of putative regulatory PD SNPs (225 or 88.6%) were located in distal accessible enhancers region, either within introns or intergenic regions (Figure 2G).

To test the accuracy of our regulatory variant prediction, we focused on 54 of the predicted putative regulatory PD SNPs that were found to be heterozygous in the TH-Rep1 cell line where our ATAC-seq data was derived from, thereby allowing allelic imbalance analysis of their chromatin accessibility. Using a chi-squared test to compare the number of reads aligning either to the reference allele or the PD-associated allele at each of the 54 variants, we found 46 out of 54 loci (or 85.2%) to show significant allelic imbalance in at least one of the measured time points of differentiation, with 40 loci (or 74%) remaining significant after multiple testing correction (Figure 2H). This indicates a strong predictive power of our approach.

Distribution and PD-GWAS significance of the of the 254 putative regulatory PD SNPs are depicted in Figure 2I. As expected, most frequent and most significant PD-associations were localized on chromosomes 4 and 17 (23 and 206, respectively). However, many significant associations were also found at other chromosomes, with regulatory PD SNPs present across a total of 10 different chromosomes (Figure 2I, Supplementary Table S1).

For each of the putative regulatory PD SNPs we predicted one or multiple TFs whose binding is affected by the presence of the PD SNP. To further prioritize the TF-enhancer-gene interactions to those that are most likely to occur in mDANs or during their differentiation, we leveraged our time series transcriptomic data to calculate the Pearson correlation for expression (RPKM) of each TF-target gene pair across the time points (Supplementary Table S2). This yielded the most promising TF-target gene pairs with absolute correlation coefficient of at least 0.5. Based on the remaining enhancer-target gene interactions, 16 of the putative regulatory PD SNPs are associated with *Microtubule-associated protein Tau* (*MAPT)* on chromosome 17 and 5 with *alpha-synuclein* (*SNCA*) on chromosome 4, representing novel putative regulatory PD SNPs affecting these key PD genes. With a total of 57 PD SNPs linked to 24 target genes (Figure 2E, Supplementary Table S2).

### The PD-associated allele of rs1465922 can alter *SCARB2* expression

The TF-target gene pair with strongest correlation was formed by NR2C2 and SCARB2 (r=0.98). Specifically, rs1465922 was found to localize in accessible chromatin in mDANs at the *SCARB2* promoter, with the PD-associated effect allele “A” (chr4:76213717) introducing a NR2C2 binding site (Figure 3A). *SCARB2* expression is significantly induced during mDAN differentiation, with parallel increasing expression of *NR2C2* (Figure 3B). Importantly, the adjacent *FAM47E* gene shows no expression despite close vicinity of the two promoters. Thus, rs1465922 is a strong candidate for regulatory PD SNP capable of influencing *SCARB2* expression through formation of a NR2C2 binding site.

**Figure 3:**
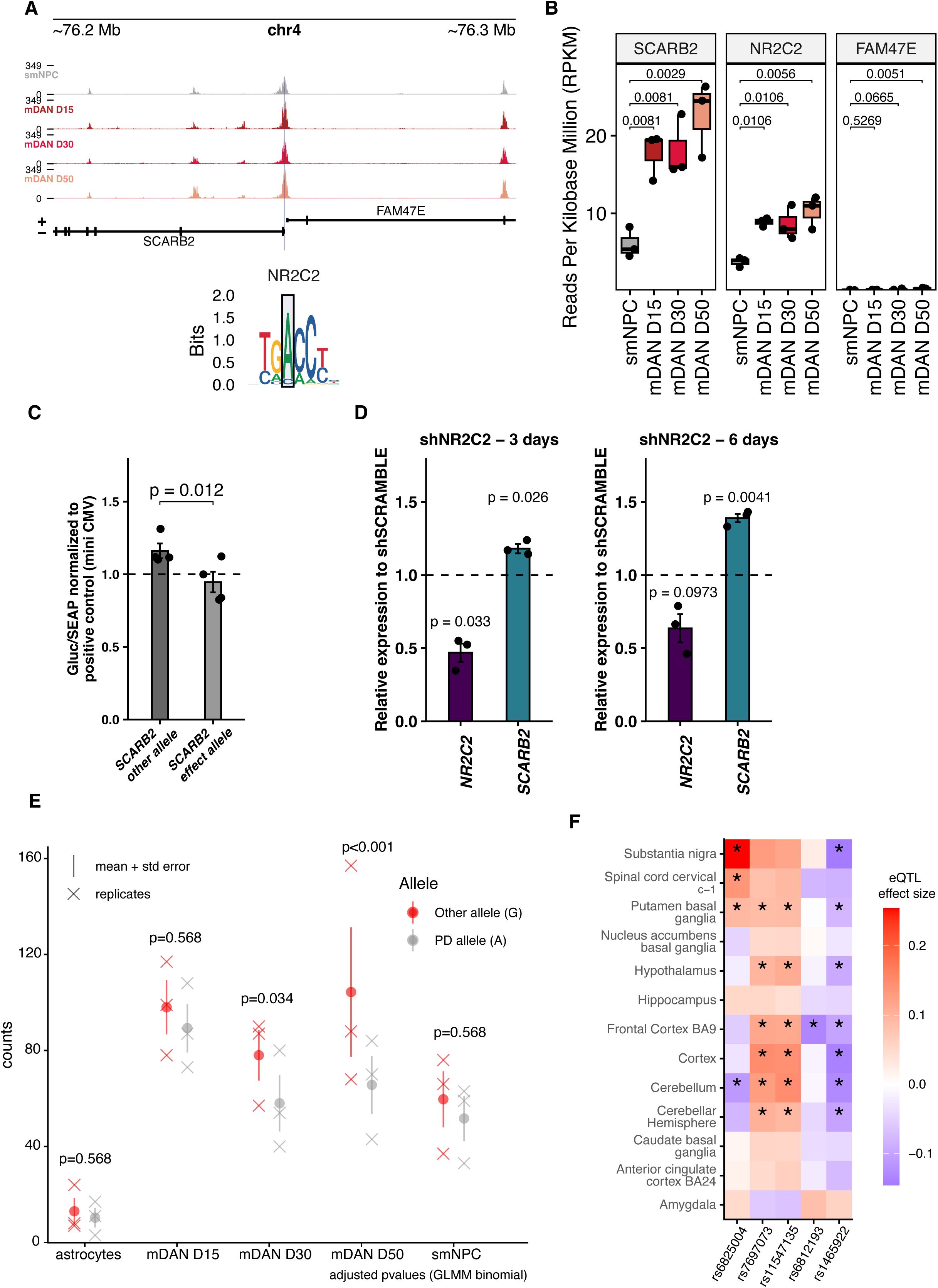
The PD-associated allele of rs1465922 modulates *SCARB2* expression. **A)** Overview showing the enrichment of open chromatin in smNPCs (grey), mDAN D15 (dark red), mDAN D30 (red) and mDAN D50 (orange) at the *SCARB2* locus The PD-associated allele (A) are highlighted in grey in the corresponding transcription factor motifs. **B)** *SCARB2*, *NR2C2* and *FAM47E* levels measured across the differentiation by RNA-seq in smNPCs and mDAN neurons. The values represent FDR of expression change compared to smNPCs. **C)** Ratio of luciferase levels over its internal control (secreted alkaline phophatase - SEAP) normalized to positive control reporter in TH-Rep1 differentiated neurons at day 11 for *SCARB2*. Bars represent mean of four biological replicates ± SD. Data points represents individual biological replicates. The statistical significance for luciferase measurements of effect allele transduced cells against other allele transduced cells was assessed by a two-sample t-test. **D)** Relative mean expression levels (± SD) of *SCARB2* and *NR2C2* following NR2C2 knockdown in TH-Rep1 differentiated neurons. The statistical significance for RT-qPCR measurements of knockdown cells against cells tranduced with a siSCRAMBLE was assessed by a two-sample t-test. **E)** Reads mapping to the effect (grey) or the non-effect allele (red) in 3 independent replicates in smNPCs, mDAN D15, mDAN D30, mDAN D50 and astrocytes. Generalized linear mixed model (random effect for the replicate) using the binomial distribution was performed to assess statistical significance between the effect allele and the non-effect allele, p-value correction was done using the Benjamini-Hochberg method (mDAN D15, *P* = 0.568; mDAN D30, *P* = 0.043; mDAN D50, *P* = 8e^−05^). **F)** eQTL effect size from GTEX v8 data for rs1465922, rs7697073, rs11547135, rs6812193 and rs1465922. The asterisk represents the p-value of a t-test comparing the computed eQTL effect size to eQTL effect size of 0.

The expression patterns of *SCARB2*, *NR2C2*, and *FAM47E* were confirmed by RNA-seq analysis of an independent TH-mCherry reporter cell line (Supplementary Figure S4A) (22) and in additional 95 iPSC lines differentiated into mDANs and profiled by the FOUNDIN-PD consortium (Supplementary Figure S4B) (44).

Importantly, the SCARB2 locus shows the presence of active chromatin modifications in ChIP-seq analysis of human substantia nigra samples from the International Human Epigenome Consortium (IHEC) (Supplementary Figure S4C). This includes histone H3 lysine 36 trimethylation (H3K36me3) in the gene bodies, consistent with active transcription (45) (46) (47). At the same time, repressive histone modifications such as lysine 9 and lysine 27 trimethylation of histone H3 are absent, confirming *in vivo* relevance of the expression of the candidate target gene in human midbrain. The *in vivo* relevance was further confirmed by the analysis of published snRNA-seq data of cell types in human substantia nigra (36), that confirmed modest but reliable level of expression for both *SCARB2* and the putative regulator, *NR2C2* in human mDANs (Supplementary Figure S4D). Their expression could also be detected in many other subpopulations of neurons and glial cells such as astrocytes and microglia. Since correlation is not causation and observed expression changes during differentiation are an outcome from integration of multiple inputs, we decided to directly test the ability of the variant to alter gene expression in mDANs. We cloned sequences flanking the NR2C2 TFBS upstream of a *Gaussia* luciferase (Gluc) reporter gene with both the PD-associated effect and the non-effect allele cloned into otherwise identical lentiviral constructs. Neuron cultures undergoing differentiation to mDANs were transduced on day 9 of differentiation and signals for luciferase reporter and secreted alkaline phosphatase (SEAP) control were measured on day 11 (Figure 3C). Consistent with our predictions, the presence of the PD-associated effect allele of *SCARB2* promoter altered reporter expression, with significant downregulation observed (Figure 3C).

Next we asked whether similar downregulation of *SCARB2* could be observed also in other NR2C2 expressing cells. We selected two common cell lines, HEK293 and HepG2, with higher and lower levels of NR2C2 expression compared to mDANs, respectively (Supplementary Figure S5A). Luciferase assays with SCARB2 promoter constructs in both cell lines showed a trend of reduced expression in the presence of PD-associated effect allele, but only in HepG2 cells this reduction was found to be significant (Supplementary Figure S5B-C). To investigate whether NR2C2 can directly bind the *SCARB2* locus, we analyzed existing NR2C2 ChIP-seq data from the ENCODE project (5). Interestingly, two of the cell lines (K562 and WTC11) showed no NR2C2 binding at *SCARB2* locus, while HepG2 cells had multiple binding sites, including at the promoter (Supplementary Figure S5D), indicating that NR2C2 can interact with *SCARB2* locus.

To further test the role of NR2C2 on *SCARB2* expression, we used lentiviral shRNA delivery to knock-down NR2C2 starting from day 9 of mDAN differentiation (Figure 3D). This significantly reduced *NR2C2* expression > 50% at three days post-transduction and was accompanied by a significant increase in *SCARB2* expression (Figure 3D). *SCARB2* expression was further increased at 6 days post-transduction, consistent with the results of the reporter gene assays. This result suggests that our experimental cell line (TH-Rep1) carries the PD associated allele mediating NR2C2 binding. Therefore, we analysed whole genome sequencing data of the mCherry-reporter cell line used in the experiments and confirmed that the cell line was heterozygous for rs1465922 at the *SCARB2* locus, carrying both alleles, capable of forming an NR2C2 binding site at the PD-associated allele, in keeping with our knock-down experiments. To confirm the result in an independent cell line with a different genotype, we carried out NR2C2 depletion in our second TH-mCherry reporter cell line (TH-Rep2) which is homozygous for the PD effect allele of rs1465922 (Supplementary Figure S5E). This resulted in even higher *SCARB2* induction already at 2 days post-transduction, confirming a regulatory interaction between NR2C2 and *SCARB2* gene. The induction at 6 days post-transduction was not found to be significant.

Since the TH-Rep1 cell line used for the time series ATAC-seq analysis was heterozygous for rs1465922 at *SCARB2* locus, we set out to test whether the PD-associated effect allele was sufficient to introduce allelic imbalance at the level of chromatin accessibility. No significant difference could be detected for the number of reads mapping to either the PD-associated effect allele or the non-effect allele in iPSC-derived smNPCs or astrocytes (Figure 3E). The overall number of mapped reads in astrocytes was also markedly lower. However, the read numbers were 4-5 times higher in the neuronal cells showing high expression of *SCARB2*, with a significant allelic imbalance in read numbers detectable specifically in mDANs. In keeping with the reporter gene assays and knock-down experiments, the presence of the PD-associated effect allele was accompanied by lower chromatin accessibility, indicating lower expression. This is particularly interesting given that recent functional screening for proteins required for lysosomal glucocerebrosidase (GCase) activity in neurons identified SCARB2 as one of the key proteins required for lysosomal homeostasis, a process dyregulated in PD (48).

This prompted us to ask whether rs1465922 is associated with reduced *SCARB2* expression also *in vivo*. To address this question, we studied the expression quantitative trait loci (eQTL) data for human brain regions from the Genotype-Tissue Expression (GTEx) project (49). Figure 3F shows the eQTL effect size for five genetic variants at the *SCARB2* locus, including variants rs6812193 and rs6825004 associated to PD, rs7697073 associated with REM-sleep behaviour disorder (RBD), and rs11547135 associated to viral infection, in addition to our SNP of interest, rs1465922 (50) (51) (52). rs6812193 did not show any significant associations with *SCARB2* expression in most of the brain regions, while rs7697073 and rs11547135 were significantly associated with increased *SCARB2* expression in several brain regions (Figure 3F). The expression association of rs6825004 varied between brain regions, with increased expression observed in particular in the substantia nigra. In contrast, rs1465922 was significantly associated only with decreased expression of *SCARB2*, with largest effect size observed in the substantia nigra. This is in line with reduced transcription and chromatin accessibility observed for this allele in gene reporter assays and ATAC-seq analysis in mDANs, respectively, and with higher expression upon NR2C2 depletion. Altogether, indicating that rs1465922 is a regulatory PD SNP contributing to the expression of SCARB2 in human mDANs.

### PD-associated allele of rs144814361 modulates *BAG3* promoter activity

Next we wanted to test whether another putative regulatory PD SNP could be validated following similar framework. Often rare variants show bigger effect sizes in genetic studies, and while the current study focuses on common variants, they still show considerable differences in their minor allele frequencies. Among our shortlisted candidates (Figure 2I, Supplementary Tables S1-S2), we identified rs144814361 at the *BAG3* locus as the SNP with lowest minor allele frequency (MAF = 0.00255102). rs144814361 is the lead variant at the *BAG3* locus, with only one other variant passing the PD association threshold of p = 5×10^−8^ and not found to be in linkage disequilibrium (LD) with other PD SNPs (Figure 2I). rs144814361 is located at the *BAG3* promoter within accessible chromatin in mDANs where the PD-associated effect allele “T” (chr10:119651405) is predicted to create a strong binding site for LHX1 and other LIM-homeodomain family (HD-LIM) TFs (Figure 4A). Indeed, the 19 different binding motifs of eight different HD-LIM family TFs resemble closely the binding site created by the variant (Supplementary Table S3), while the predicted binding energy change is most significant for LHX1 (p = 0.005867). *BAG3* and *LHX1* expression levels show negative Pearson correlation (r=-0.55), with highest *BAG3* expression in smNPCs, reduced during mDAN differentiation, and increased again after 50 days of differentiation while *LHX1* shows an opposite behaviour (Figure 4B).

**Figure 4.**
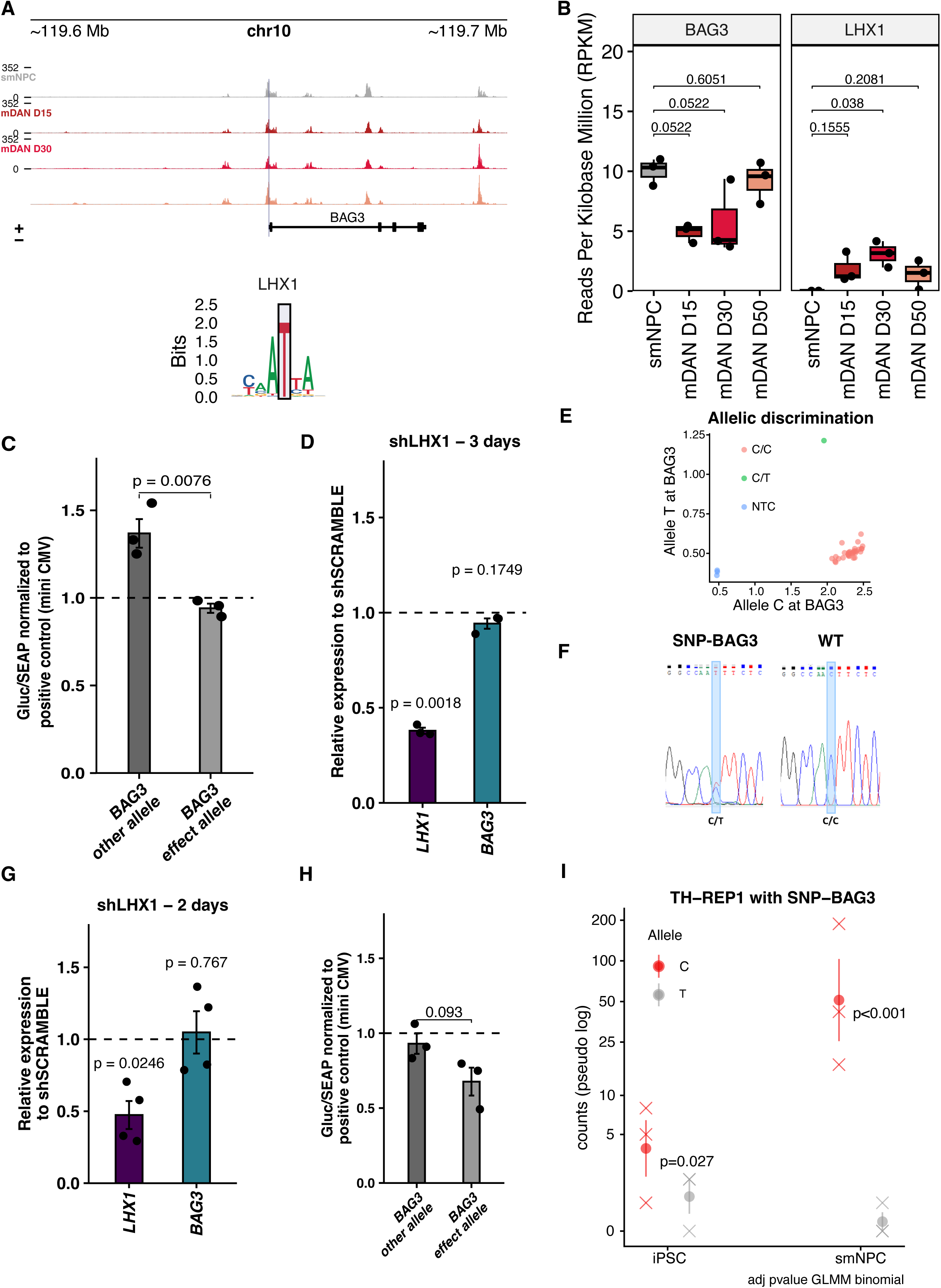
PD-associated allele of rs144814361 modulates *BAG3* promoter activity. **A)** Overview showing the enrichment of open chromatin in smNPCs (grey), mDAN D15 (dark red), mDAN D30 (red) and mDAN D50 (orange) at the *BAG3* locus. The PD-associated allele (T) is highlighted in grey in the corresponding transcription factor motif. **B)** *BAG3* and *LHX1* levels measured across the differentiation by RNA-seq in smNPCs and mDAN neurons. The values represent FDR of expression change compared to smNPCs. **C)** Ratio of luciferase levels over its internal control (secreted alkaline phophatase - SEAP) normalized to positive control reporter in TH-Rep1 differentiated neurons at day 11 for *BAG3*. Bars represent mean of four biological replicates ± SD. Data points represents individual biological replicates. The statistical significance for luciferase measurements of effect allele transduced cells against other allele transduced cells was assessed by a two-sample t-test. **D)** Relative mean expression levels (± SD) of *BAG3* and *LHX1* following LHX1 knockdown in TH-Rep1 differentiated neurons. The statistical significance for RT-qPCR measurements of knockdown cells against cells tranduced with a siSCRAMBLE was assessed by a two-sample t-test. **E)** The presence of the BAG3 SNP variant was assessed in 34 iPSC clones using human TaqMan® SNP Genotyping Assays. One clone REP1-SNP-BAG3-TH-mcherry (named TH-Rep1-SNP-BAG3) is heterozygous for SNP BAG3 (C/T). **F)** Sanger sequencing of the prime edited SNP-BAG3 clone and the WT TH-Rep1 cell line. **G)** Relative expression levels of *BAG3* and *LHX1* following LHX1 knockdown in TH-Rep1-SNP-BAG3 differentiated neurons. The statistical significance for RT-qPCR measurements of knockdown cells against cells tranduced with a siSCRAMBLE was assessed by a two-sample t-test. **H)** Ratio of luciferase levels over its internal control (secreted alkaline phophatase - SEAP) normalized to positive control reporter in smNPCs for *BAG3*. Bars represent mean of three biological replicates ± SD. Data points represents individual biological replicates. The statistical significance for luciferase measurements of effect allele transduced cells against other allele transduced cells was assessed by a two-sample t-test. I) Reads mapping to the effect (grey) or the non-effect allele (red) in iPSCs and smNPCs derived from TH-Rep1 SNP-BAG3 cell line. A generalized linear mixed model (random effect for the replicate) using the binomial distribution was performed: fixed effect on allele counts (iPSC, *P* = 0.027; smNPC, *P* = 7.7e^−08^).

Similarly to *SCARB2*, *BAG3* was expressed in progenitors and mDANs derived also from other iPSC lines (Supplementary Figure S6A-B), and the locus showed a chromatin state consistent with active transcription in human substantia nigra samples (Supplementary Figure S6C). However, based on snRNA-seq analysis of human substantia nigra, most BAG3 expression *in vivo* occurs in astrocytes, with only some subpopulations of mDNAs and other neurons showing BAG3 expression (Supplementary Figure S4D). Focusing on the putative regulators of *BAG3*, most HD-LIM TFs became induced during mDAN differentiation, with the exception of *LHX6* and *ISL2*, that were either absent or expressed only in smNPCs, respectively (Supplementary Figure S6D). According to *in vivo* snRNA-seq data, HD-LIM TF expression was largely limited to different neuronal population, with only LHX1, LMX1A, and LMX1B detected in mature mDANs (Supplementary Figure S6E).

Lentiviral delivery of *Gaussia* luciferase (Gluc) reporter gene constructs carrying either the PD-associated effect allele creating the TFBS or the otherwise identical non-effect allele into 9 days differentiated mDANs resulted in a significant downregulation in the presence of the PD-associated allele (Figure 4C). However, the depletion of LHX1 did not alter the expression level of *BAG3* at 3 days after shRNA transduction (Figure 4D).

To better understand why only *SCARB2* expression was altered by the depletion of the predicted TF regulator, although also the *BAG3* alleles showed differential expression in reporter gene assays, we analysed again the whole genome sequencing data of the mCherry-reporter cell line (TH-Rep1) used in the experiments. Importantly, consistent with the low minor allele frequency of the PD-associated allele, the cell line was found to be homozygous for the non-effect allele (“C” at the position of rs144814361 at the *BAG3* locus), rendering the cell line devoid of LHX1 (or other HD-LIM TF) binding site and providing a potential explanation for the lack of change in *BAG3* expression.

In order to confirm the predicted impact of the PD-associated allele of rs144814361 on BAG3 promoter, we proceeded with genome editing of the TH-Rep1 cell line by using prime editing to insert the “T” allele at the position chr10:119651405 in the *BAG3* promoter (Supplementary Figure S7). The prime-editing strategy used to introduce the variant in iPSCs included the design of the pegRNA and a PE3 nicking sgRNA, together with the genomic context of the targeted *BAG3* promoter region (24,53). Following isolation, expansion, and selection by SNP genotyping assay (Figure 4E), one edited clone (1/34) was validated by Sanger sequencing (Figure 4F).

The pluripotency of the edited iPSC clone, designated TH-Rep1-SNP-BAG3, was confirmed by testing the expression of known marker genes by PCR and immunocytochemistry (Supplementary Figure S8A-B). The iPSCs were successfully induced into neural progenitor cells (NPCs) (Supplementary Figure S8C-D), and subsequently differentiated into mDANs with an efficiency comparable to the wildtype (WT) parental reporter cell line, as confirmed by mCherry signal (Supplementary Figure S8E-F).

Using the genome edited line we again performed a LHX1 knockdown in differentiating mDANs as described above (Figure 4G). However, no change in *BAG3* expression could be detected, suggesting that rs144814361 may not act as a regulatory variant, or that LHX1 is not the only TF binding to the created site. To address this we transduced the reporter gene constructs carrying the two BAG3 alleles into smNPCs that do not express LHX1 but do express BAG3 (Figure 4B). Interestingly, the reporter carrying PD-associated effect allele did show a trend of reduced expression also in smNPCs (p=0.093). This suggests that the effect allele could influence gene expression independent of LHX1. To directly confirm the differential regulatory activity of the PD-associated effect allele in the genome edited cells, we performed ATAC-seq analysis of iPSCs and smNPCs derived from the TH-Rep-SNP-BAG3 cell line. Consistent with low expression of *BAG3* in iPSCs compared to smNPCs, the chromatin accessibility at the *BAG3* locus was 10-fold lower in iPSCs (Figure 4I). Despite the low number of sequencing counts, the PD-associated effect allele showed a significantly lower accessibility even in iPSCs. And importantly, in smNPCs this difference was markedly stronger and highly significant, with most ATAC-seq reads mapping to the major “C” allele. Indicating that PD-associated minor allele of rs144814361 can significantly modulate *BAG3* promoter activity, possibly by creating a binding site for several HD-LIM family TFs.

## Discussion

GWAS studies have demonstrated disease–and trait-associated variants to be mostly located into the non-coding genome (54,55). The introduction of variants in *cis*- and *trans-*regulatory regions can influence the phenotype of a cell by altering the binding of TFs and their target gene expression in a cell type-specific manner. Thus, with the majority of disease associated variants located in cell type-specific regulatory regions of the genome, the main challenge is to reliably predict their target genes and their regulatory impact in a cell type. To overcome this challenge, we have taken an integrative multiomics approach. Since PD-associated variants were found to be enriched in the vicinity of genes expressed in mDANs, we combined gene expression, chromatin accessibility, and LowC chromosome conformation data to map target genes and their interactions with regulatory regions in human mDANs. By integrating these data with information about PD-associated variants, we computed the expected changes in binding affinities at TFBS due to the presence of the variants and predicted the affected TFs and their target genes in smNPCs and mDANs. Through this approach, we reduced 3464 PD-associated candidate variants with genome-wide significance in PD GWAS to 254 variants with potential to act as regulatory variants in mDANs and during their differentiation. Previous analysis have already studied PD-associated variants in regulatory regions in tissues (56) and major cell types of the brain such as excitatory and inhibitory neurons, oligodendrocytes, and microglia (57,58), including single nuclei analysis of mDANs (59). Moreover, detailed function of some individual variants have been studied in neuronal and microglia cultures, or in common cell lines (16,60,61). However, detailed prediction and validation of the context-specific regulatory impact is often difficult due to limited number of mDANs in post-mortem samples or lack of relevant experimental model system. The number of putative regulatory PD variants identified in our analysis (254) is comparable to the 165 dysregulated PD genes identified previously in single nuclei analysis of mDANs from post-mortem brains (59). The enhancer-target gene prediction based on our ATAC-seq and LowC analysis indicated that predicted variants were associated with hundreds of target genes in mDANs, in addition to many specific interactions in smNPCs. As expected, these genes were enriched for phenotypes like PD. As two interesting examples responsible for creating a TFBS in human mDNAs, we identified variants rs144814361 and rs1465922 at *BAG3* and *SCARB2* loci, respectively. The PD-associated allele at *BAG3* locus was predicted to create a binding site for HD-LIM family TFs, in particular LHX1 that showed a negative correlation with *BAG3* expression, while the predicted regulator of *SCARB2* expression, NR2C2, showed a strong positive correlation with its target gene. However, functional analysis of the PD-associated alleles revealed a role in downregulation of the associated target genes for both variants.

BAG3 has been shown to be involved in the development of the central nervous system in rats. Specifically, in the prenatal forebrain the expression of BAG3 can be detected in the medial telencephalic wall of the lateral ventricle and in the Cajal-Tetzius cells of the developing cortex. In the postnatal brain the expression of BAG3 increased in the neurons of the cortex and hippocampus, followed by a major decrease, indicating an important role for dynamic BAG3 expression during neurodevelopment (62). Moreover, as we also show, BAG3 is found in glial cells of the midbrain. Interestingly, BAG3 shows a higher expression in the hippocampus of one year old mice, suggesting that BAG3 may be implicated in protein homeostasis and clearance during aging, consistent with a misregulation of this pathway in age-related diseases such as PD (63). In addition, expression of BAG3 was detected in rat dorsal midline glia cells in the midbrain and in the spinal cord but not in the ventral midline of the same brain tissues (64).

More relevant for PD, increased expression of BAG3 was reported in transgenic mice overexpressing mutated human SNCA in dopaminergic neurons (65). Additionally, accumulation of SNCA was detected in BAG3-depleted cells while overexpression of BAG3 produced the opposite effect through increased autophagy (65). Finally, regulatory variation at BAG3 locus has been recently associated to PD as PD-associated SNPs were found to coincide with regulatory regions in microglia as defined by accessible chromatin and acetylation of histone H3 at lysine 27 (60). In our study, we also found a PD-associated SNP (rs144814361) in open chromatin at the BAG3 promoter in smNPCs and mDANs. However, the strength of our integrative computational approach was to also use chromatin conformation data, together with prediction of TF binding change at the accessible site, to allow more precise prediction. Moreover, our reporter gene assays showed that the presence of rs144814361 results in altered gene expression in smNPCs and mDANs. Finally, use of genome editing to generate a isogenic cell line carrying the PD-associated variant confirmed its regulatory potential through strong modulation of promoter chromatin accessibility.

LHX1 was predicted as the most likely TF to bind to the PD-associated BAG3 allele. It is a transcription factor expressed in postnatal and adult mouse and rat Purkinje cells (66,67) that are responsible for body’s motor functions. In addition, LHX1 plays a role in the control of motor neuron migration (68). A common limitation of studies on regulatory variation is the difficulty to identify the exact TF or TFs whose binding is affected by the variant. In keeping with this, also LHX1 belongs to the HD-LIM family of TFs, that are characterized by a LIM interaction domain and a homeodomain, and act broadly as transcriptional regulators during neuronal development and function, sharing highly similar DNA-binding motifs. A lower luciferase activity was observed in both mDANs and smNPCs when the putative LHX1 binding site is present, suggesting a repressive effect mediated by several HD-LIM TFs rather than LHX1 alone. Consistent with this, introduction of the PD-associated allele by genome editing resulted in reduced chromatin accessibility across different cell states, further indicating that multiple transcription factors may contribute to regulation at this locus. These data, together with previous reports, indicate that genetic variation in *BAG3* expression regulation in several cell types could play a role in PD development. Further studies will be needed to understand modulation of BAG3 by rs144814361 in detail.

*SCARB2* locus has been associated with rapid-eye movement (REM) sleep behavior disorder (RBD) (52) that typically precedes PD as a prodromal stage of dopaminergic neurodegeneration years before typical motor symptoms become apparent (69–72). SCARB2 is the transporter of the enzyme GCase encoded by the glycosylcerebrosidase (GBA) gene, mutations in which are the most common genetic risk factor for PD (73). Consistently, depletion of SCARB2 expression was shown to be detrimental for lysosomal GCase activity in neurons (48). Several SNPs in the proximity of SCARB2 locus have been associated to PD through GWAS. For example, the SNPs rs6812193 and rs6825004 have been associated to PD in populations from European ancestry (74). However, these alleles/variants had no effect on SCARB2 expression at the transcriptional or protein levels (51), highlighting the difficulty of associating individual SNPs to their respective target genes identified via GWAS. More recently, a non-coding variant rs11547135, was independently associated with both PD and SCARB2 expression in blood and brain tissue, in keeping with the eQTL results in Figure 3F (75). In our study, we overcame some of the challenges by associating prediction from TF binding change with chromatin conformation and accessibility data, narrowing the analysis to the study of rs1465922, and its ability to introduce a putative NR2C2 TFBS in the promoter of *SCARB2*.

NR2C2, also known as testicular receptor 4 (TR4), is an orphan nuclear receptor. In mouse, it has been shown that NR2C2 knockout leads to difficulties in motor coordination (76). In rats, Young *et al*. have shown that NR2C2 is highly expressed in the brain, the spinal cord, the ganglia and neuronal epithelia (77). Also, at the embryonic stage in rats, NR2C2 is abundant in the central nervous system (CNS), as well as in the cerebellum and the hippocampus in postnatal rats (78). In our study, the creation of a NR2C2 TFBS through the presence of rs1465922 led to a decrease in luciferase activity in a *SCARB2* promoter reporter assay, indicating a possible repressive action of NR2C2 on *SCARB2* expression. In contrast, knockdown of NR2C2 led to an increase in expression of *SCARB2* in human neurons, together highlighting NR2C2 as a repressor of *SCARB2*.

Finally, beyond the results on regulatory variants, our analysis of 3D chromatin conformation indicate that differentiation into mDANs is accompanied by a pronounced reorganization of higher-order genome architecture, characterized by increase in heterochromatin-like regions, strengthening of euchromatic interactions, and large-scale merging of TADs. The emergence of fewer but larger and more strongly insulated TADs in mDANs is consistent with a neuronal chromatin state in which local regulatory structures are simplified, potentially reflecting altered CTCF occupancy and compartmentalization (38). Notably, these architectural changes occur largely without detectable genome-wide alterations in transcription or chromatin accessibility at the 500 kb scale, suggesting that 3D genome reorganization during mDAN differentiation may primarily shape the regulatory landscape by constraining long-range interactions rather than directly driving bulk transcriptional changes. A more detailed analysis of the identified long-distance regulatory interactions specific to mDANs, and the possible impact of PD-associated variants on their activity, will be important for improved understanding of PD genetics and disease-risk.

## Conclusions

We provide a resource of putative candidates of regulatory PD SNPs – and a framework for their identification and functional validation – and experimentally validate the regulatory potential of two of the variants. This resource will facilitate future studies that will be needed to test their role as regulatory variants, and to confirm their contribution to gene regulation and development of PD. With the overall goal to allow cell type-specific predictions of disease-associated gene network alterations from personal genomes, and generation of improved risk scores incorporating regulatory impact of disease-associated variants.

## Supplementary data

Supplementary data is available online.

## Acknowledgments

The sequencing was performed at the Genomics Platform of Luxembourg Centre for System Biomedicine. The computational analysis presented in this paper were carried out using the HPC facilities of the University of Luxembourg. All schematic representations were created in BioRender using license Catillon, M. (2025) https://BioRender.com/5kzs36o.

## Authors Contributions

D.G., J.O., T.S., R.K., and L.S. conceived the project and designed the experiments. D.G. established the LowC assay, performed most of the experiments and bioinformatic analysis, interpreted data and prepared figures. J.O. and B.G.R performed experiments and bioinformatic analysis, and interpreted data. M.C. performed experiments and genome editing. J.S. and A.G performed LowC data analysis in consultation with H.K. and L.S.. N.B., D.H., and M.S. established the SNEEP pipeline and performed the analysis. A.G. performed bioinformatic analysis and prepared figures. Z.L., J.O., and P.M. performed MAGMA analysis and whole genome sequencing. A.R. designed and generated the iPSC reporter lines in consultation with C.K.. L.S. and R.K. supervised the project. D.G., J.O., and L.S. wrote the manuscript. All authors revised and approved the final manuscript.

## Data availability

The source high-throughput sequencing fastq files are available under restricted access at https://ega-archive.org/, under the accession numbers EGAD00001009288 and EGAD50000002258. The processed data can be visualized using UCSC genome browser at: https://genome-euro.ucsc.edu/cgi-bin/hgTracks?db=hg38&hubUrl=https://biostat2.uni.lu/PD_SCARB2/hub.txt

Analysis code and the Singularity definition file can be found in the following repository: https://github.com/sysbiolux/PD-rSNP_SCARB2

Data used in the preparation of this article were obtained on May, 22, 2023 from the Parkinson’s Progression Markers Initiative (PPMI) database (www.ppmi-info.org/access-data-specimens/download-data), RRID:SCR_006431. For up-to-date information on the study, visit www.ppmi-info.org.

## Declarations

### Ethics approval

Ethical approval for experiments with human iPSC-derived cell types was obtained from the Ethics Review Panel of University of Luxembourg (Reference ERP 25-031 EpiCNS).

