## Supplementary Data File for "Identification of Parkinson’s disease-associated regulatory variants in human dopaminergic neurons reveals modulators of *SCARB2* and *BAG3* expression"

This pdf includes:

- Supplementary Figure S1-S8 and associated legends
- Supplementary Table S1-S3 legends

Other Supplementary files include Supplementary Tables S1-S3, provided as excel files.

#### Supplementary Figure Legends

**Supplementary Figure S1. Expression of dopaminergic neuron marker genes in iPSC-derived cells.** A-G) RNA-seq expression of indicated mDAN marker genes in TH-Rep1 cells across different cell states. Boxes indicate the interquartile range with the median shown as a central line; whiskers extend to 1.5× the interquartile range. Individual data points are shown as jittered dots. N=3 independent experiments.

**Supplementary Figure S2. 3D genome organization in smNPCs and mDANs**. **A)** (Top) Line plots showing PC1 values, calculated in 500 kb genomic bins, for smNPCs (pink) or mDANs (green) in chr 11. (Bottom) Line plots showing percentage (bp/bp) of Alu (blue) or LINE (brown) elements, calculated in 500 kb genomic bins, in chr 11. **B)** Box plot showing ATAC-seq read depth in G1, G2 and G3 groups of 500 kb genomic bins. **C)** Heatmaps showing for A-A, A-B and A-B interactions. See log2-fold change (FC) profiles at the right. See Figure 1 for details. **D)** Changes in B-B interactions in chr11 are highlighted with brown triangles.

**Supplementary Figure S3. 3D genome organization in NeuN-positive and NeuN-negative cells.**

**A)** Dot plot showing PC1 values for non-neuronal ($NeuN^{neg}$) and neuronal ($NeuN^{pos}$) cells, calculated in a 500 kb resolution. Red dots denote 500 kb bins that showed a decrease in PC1 values in non-neuronal cells (PC1 of $NeuN^{neg}$ - $NeuN^{pos}$ > 0.075 in 190 regions, G1 group), while blue dots denote 500 kb bins that showed a reciprocal increase in PC1 values ($NeuN^{neg}$ - $NeuN^{pos}$ < -0.075 in 240 regions, G3 group). The rest are shown as gray dots (G2 group). **B)** Metaplots showing A-A, B-B and A-B interactions. The color code denotes interaction levels between neighboring regions (red = strong, blue = weak). A plot based on the fold change of values between non-neuronal and neuronal cells is shown at the right for intra-A or intra-B interactions. Stronger interactions in neuronal cells are shown in magenta, while weaker interactions are shown in green. **C)** Comparison of TAD lengths in non-neuronal ($NeuN^{neg}$) and neuronal ($NeuN^{pos}$) cells. The number and median length of TADs are shown at the top. Statistical significance were calculated by Student's t-test.

**Supplementary Figure S4. TH-Rep2, iPSC-derived neurons, and *in vivo* expression data for SCARB2. A)** TH-Rep2 smNPCs were differentiated and sorted for $TH^{+}$ neurons at day 15 and day 30. The mRNA levels of SCARB2 and NR2C2 were measured across differentiation by RNA-seq. The values represent FDR of expression change compared to smNPCs. **B)** FOUNDIN-PD iPSCs-derived neurons at day 25 and day 65. The mRNA levels of SCARB2 and NR2C2 were measured by RNA-seq.

The data were obtained from FOUNDIN-PD (44). **C)** Overview showing the enrichment of H3K4me1 (light pistachio green), H3K4me3 (celadon blue), H3K36me3 (steel blue), H3K9me3 (light parma) and H3K27me3 (dark parma) in human substantia nigra at the *SCARB2* locus. The data were obtained from IHEC (7). **D)** Heatmap illustrating in vivo snRNA-seq expression of SCARB2, NR2C2, BAG3, and mDAN-specific marker genes (TH, NR4A2) across astrocytes, mDANs, microglia, and non-mDANs in post-mortem human substantia nigra tissue (36). Each column represents one cell with total of 358 889 cells represented.

**Supplementary Figure S5. A)** Comparative *NR2C2* gene expression between HepG2 and HEK293T cells, and TH-Rep1 neurons at day 11 of differentiation. **B-C)** The mean ratio (± SD) of luciferase levels over its internal control (secreated alkaline phophatase - SEAP) in HepG2 (B) and HEK293T cells (C) for *SCARB2.* **D)** ChIP-seq signal of NR2C2 binding at SCARB2 locus in three studied cell lines, HepG2 K562, and WTC11. The ChIP-seq data were obtained from ENCODE (5). **E)** Relative mean expression levels (± SD) of *NR2C2* and *SCARB2* following NR2C2 knockdown in differentiated neurons derived using TH-Rep2 cell line. The statistical significance for RT-qPCR measurements of knockdown cells against cells tranduced with a shSCRAMBLE was assessed by a two-sample t-test.

**Supplementary Figure S6. TH-Rep2, iPSCs differentiated neurons, and *in vivo* expression data for BAG3. A)** TH-Rep2 smNPCs were differentiated and sorted for $TH^+$ neurons at day 15 and day 30. The mRNA levels of BAG3 and LHX1 were measured across differentiation by RNA-seq. The values represent FDR of expression change compared to smNPCs. **B)** FOUNDIN-PD iPSCs differentiated neurons at day 25 and day 65. The mRNA levels of BAG3 and LHX1 were measured by RNA-seq. The data were obtained from FOUNDIN-PD (44). **C)** Overview showing the enrichment of H3K4me1 (light pistachio green), H3K4me3 (celadon blue), H3K36me3 (steel blue), H3K9me3 (light parma) and H3K27me3 (dark parma) in human substantia nigra at the *BAG3* locus. The data were obtained from IHEC (7). **D)** Heatmap showing RNA-seq expression of the HD-LIM family of transcription factors *ISL2, LHX1, LHX2, LHX5, LHX6, LHX9, LMX1A,* and *LMX1B* in the TH-Rep1 cell line. **E)** Heatmap illustrating *in vivo* snRNA-seq expression of the HD-LIM family of transcription factors *ISL2, LHX1, LHX2, LHX5, LHX6, LHX9, LMX1A, LMX1B* across astrocytes, mDANs, microglia, and non-mDANs in post-mortem human substantia nigra tissue (36). Each column represents one cell with total of 358 889 cells represented.

**Supplementary Figure S7. Precise SNP-BAG3 editing in iPSCs using the Prime Editing system.** The Prime editing system is composed of a Cas9-nickase (H840A) fused to a reverse transcriptase (RT) which cuts only one DNA strand, a pegRNA guides the complex to the target and also contains a primer binding site (PBS), a reverse transcription template with the desired edit. The RT copies the edit sequence into the target DNA using the pegRNA as template. The specific genomic locus (rs144814361) shows the wild-type DNA sequence, the pegRNA components (sgRNA,PBS, RT template ) marks the intended edit C -> T. The Prime editing workflow includes design, cloning, delivery, clone isolation, and validation of isogenic clones. After editing, the iPSCs were induced into NPCs, then differentiated into mDANs.

**Supplementary Figure S8. Characterization of TH-Rep1-SNP-BAG3 cell lines. A)** The expression of stemness markers OCT3/4, DNMT3B and NANOG was analysed by RTqPCR in TH-Rep1 iPSCs (2 biological replicates) and TH-Rep1-SNP-BAG3 (3 biological replicates). **B)** Immunofluorescence images of edited iPSCs showed staining for OCT3/4 and SOX2, with DAPI marking nuclei. **C)** The relative gene expression (Log2 Fold change) analysis of stemness (NANOG, OCT3/4) and neural progenitor markers (NESTIN, PAX6 and SOX1) was performed by RTqPCR on NPC TH-Rep1-SNP-BAG3 and normalized to iPSC TH-Rep1-SNP-BAG3. **D)** Brightfield images showed the morphology of the NPCs (passage 5) and neurons (day 17). **E)** Differentiated mDANs (Day 16) were co-stained with neuronal antibodies TH and MAP2, nuclei were stained with DAPI. **F)** Differentiated mDANs (day 17) expressing the TH (mcherry positive cells) were analyzed by Flow cytometry. Differentiated 17608/6 (mcherry negative) and TH-Rep1 mDANs (mcherry positive) were shown.

### Supplementary Figure 1

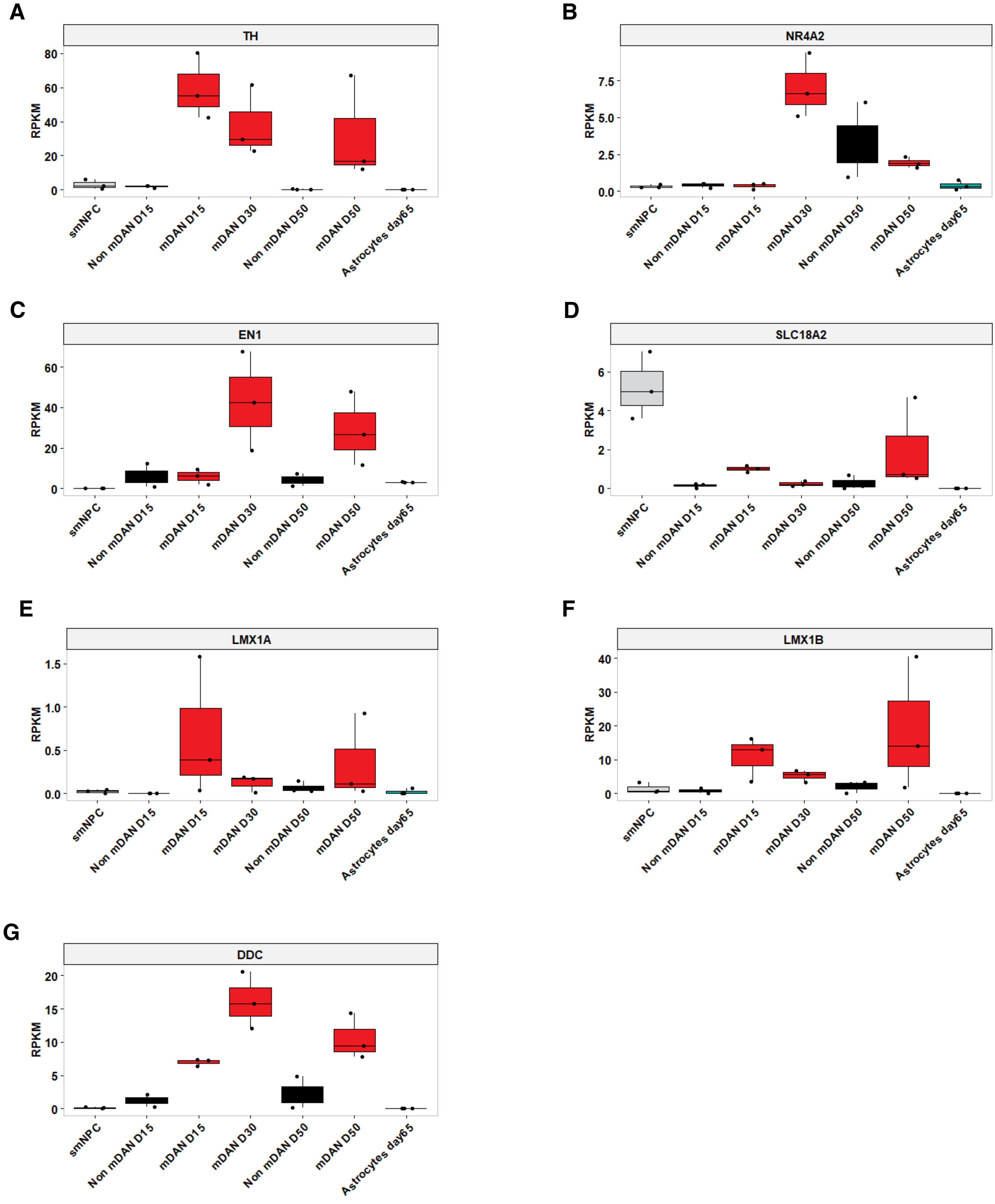

### Supplementary Figure 2

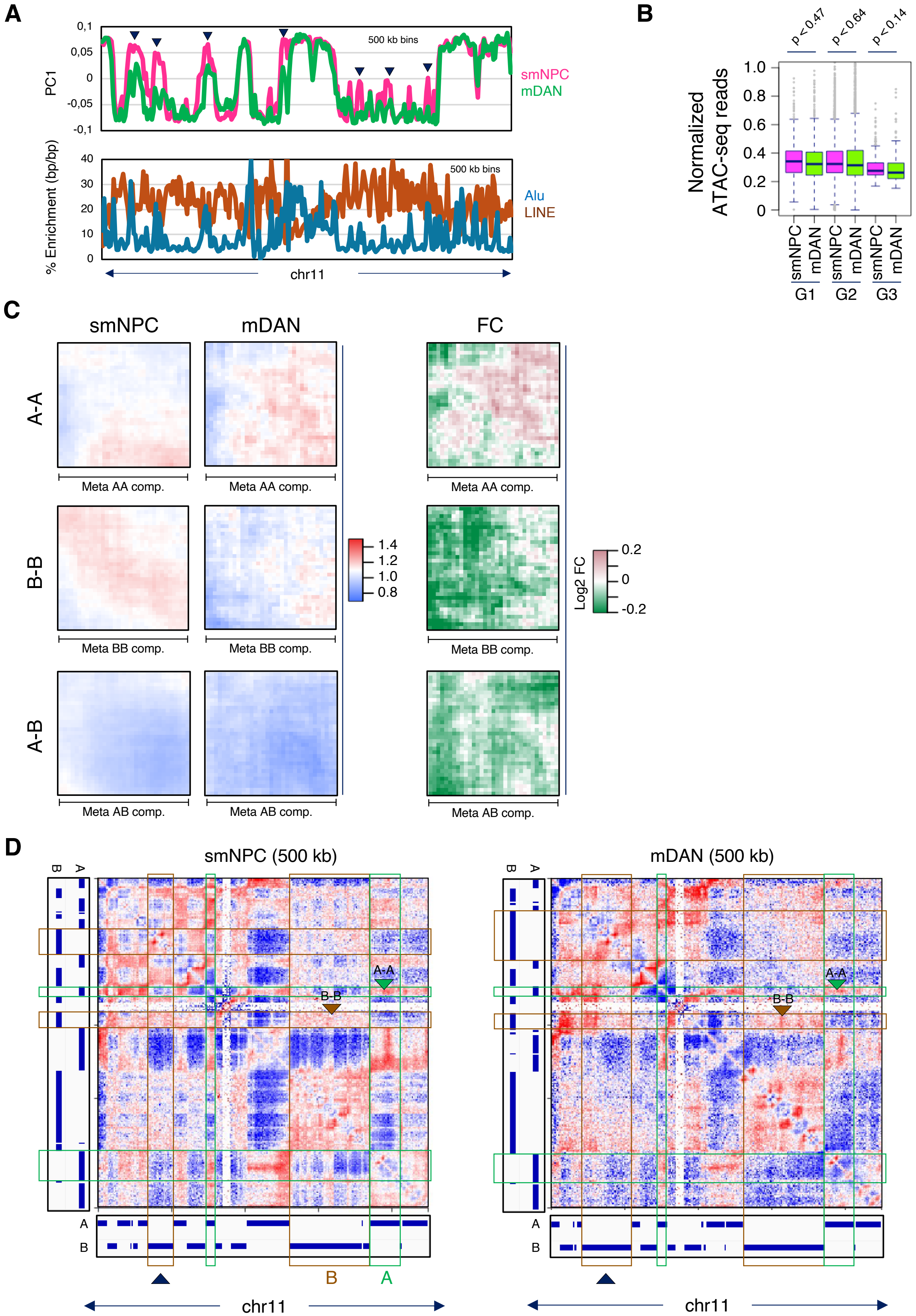

### Supplementary Figure 3

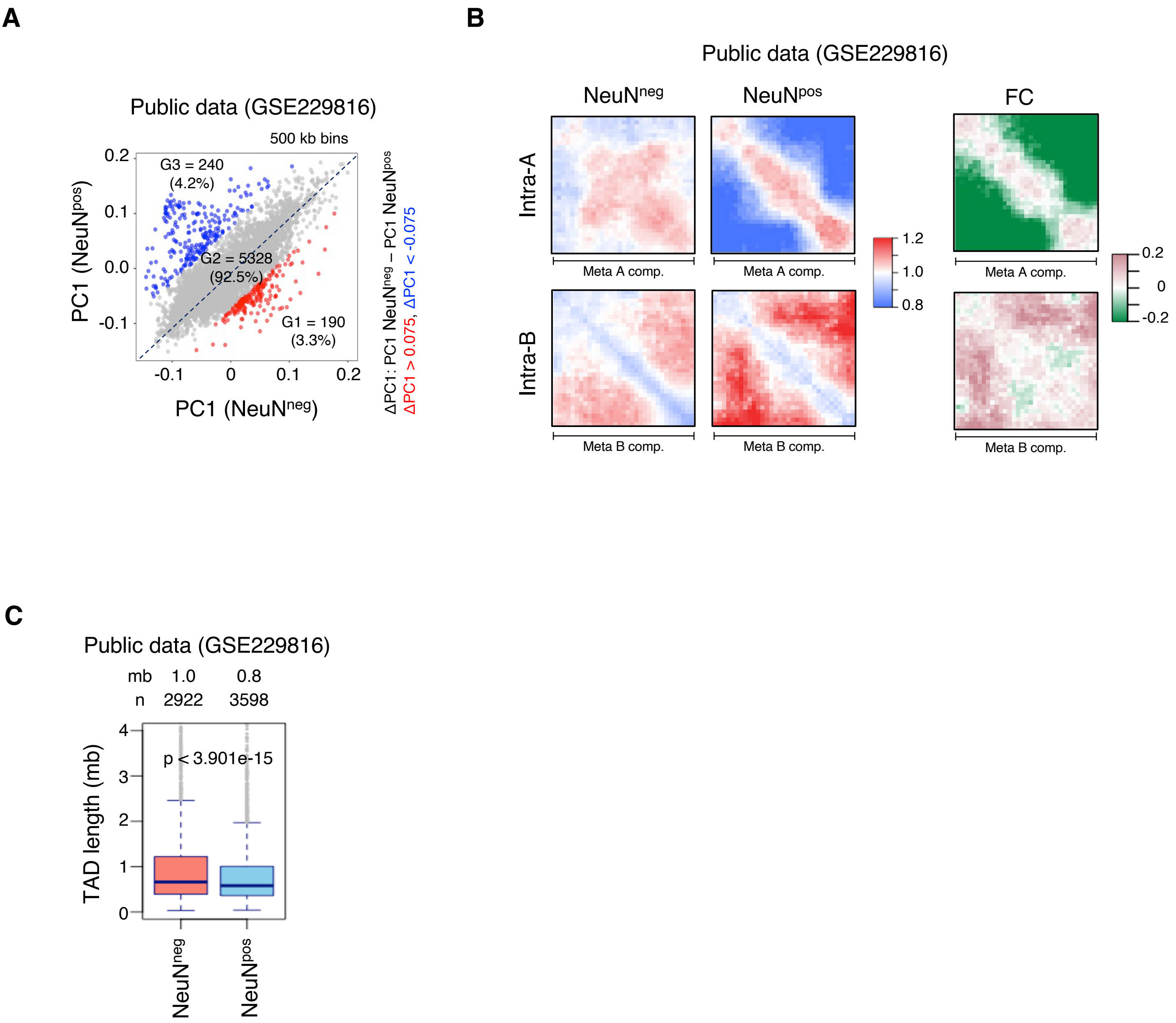

### Supplementary Figure 4

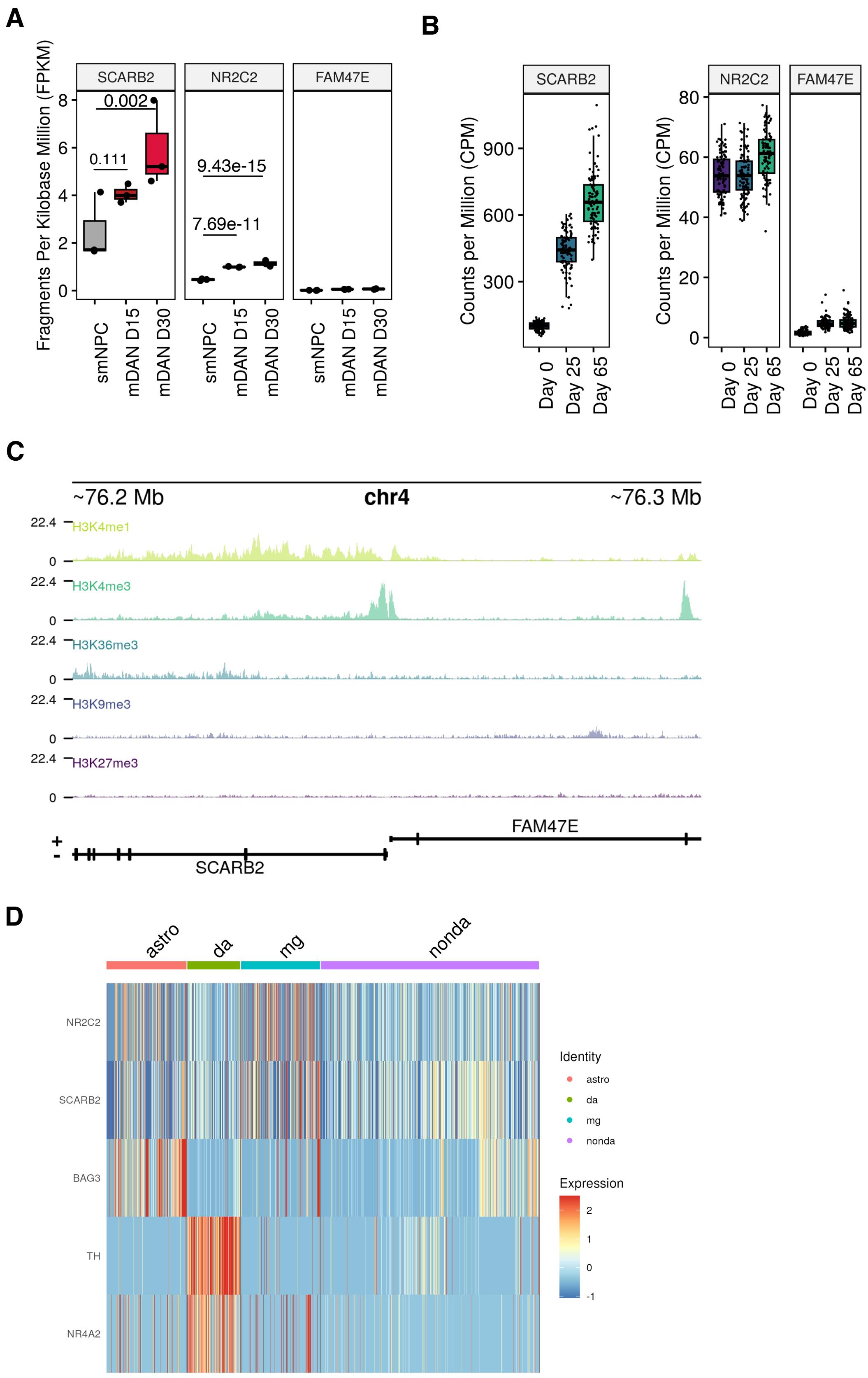

#### Supplementary Figure 5

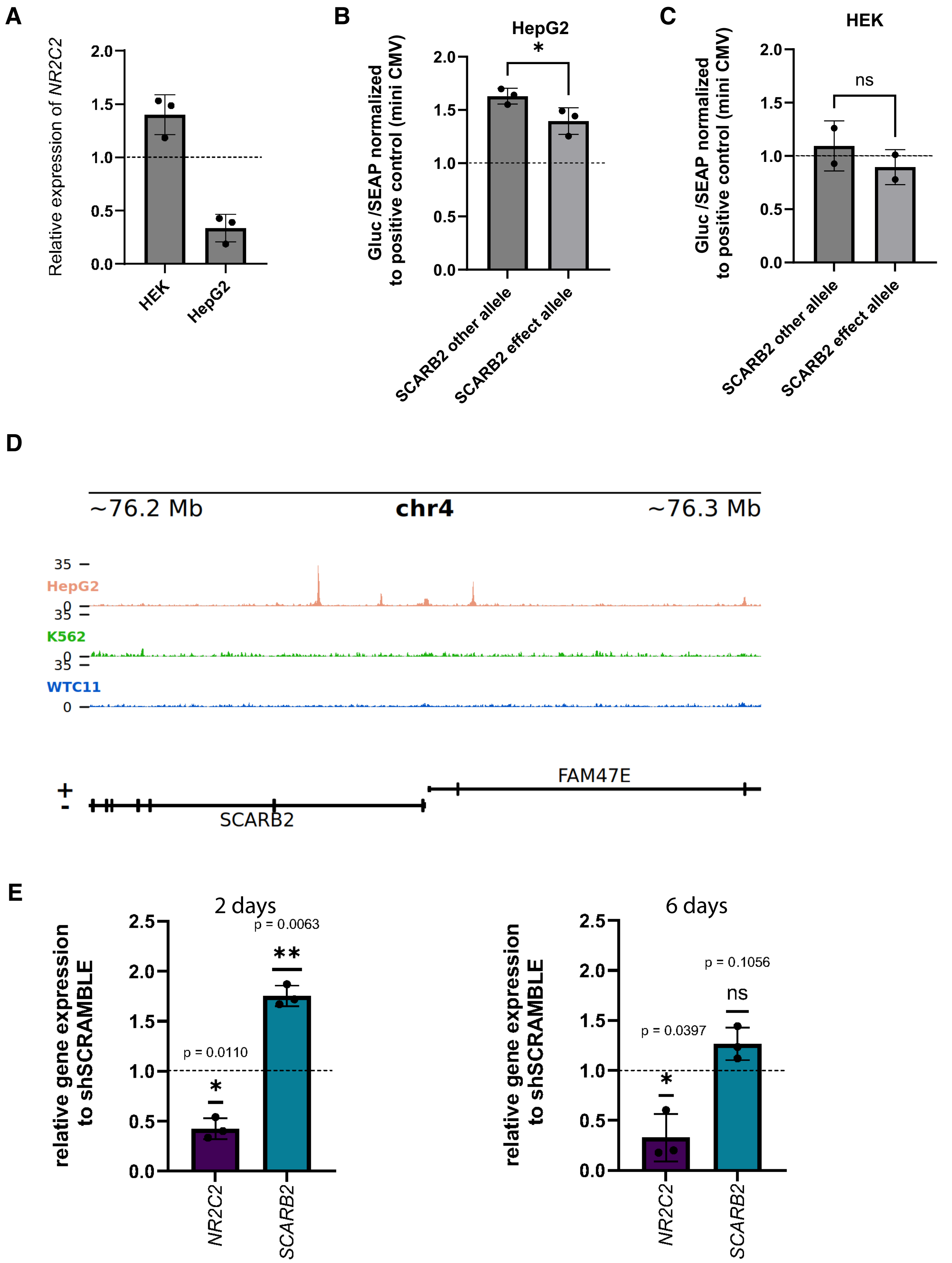

### Supplementary Figure 6

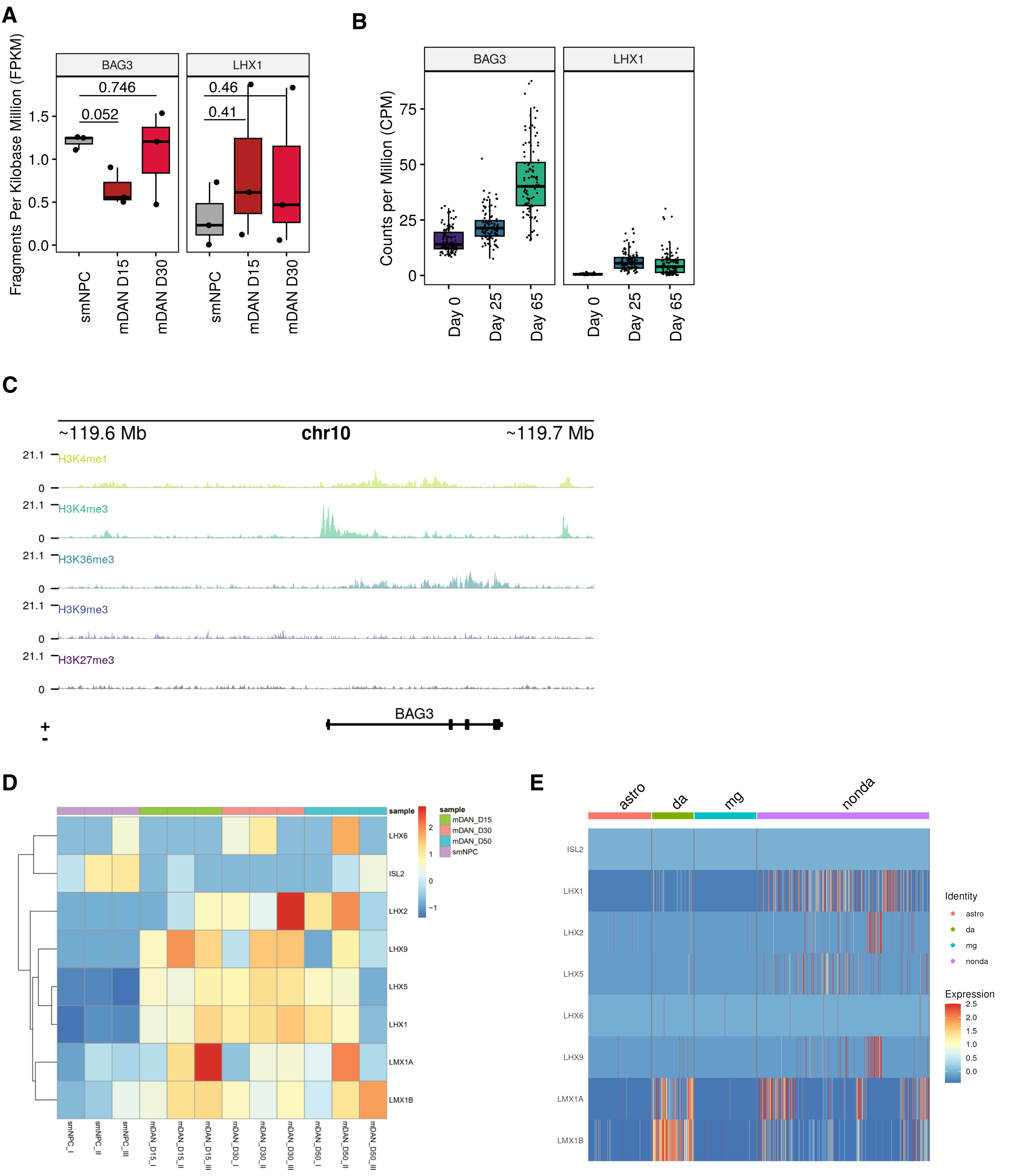

### Supplementary Figure 7

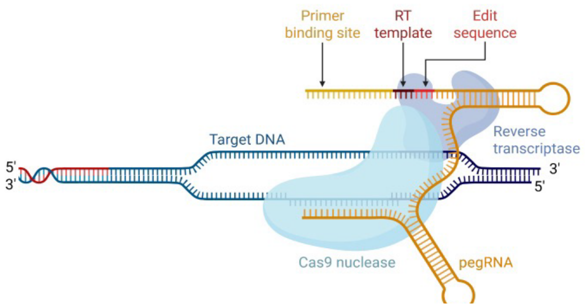


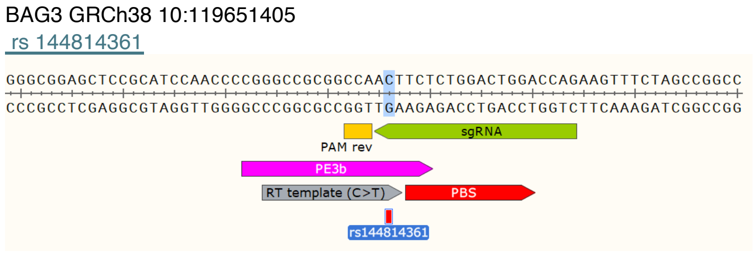


1) Cas9-nickase (H840A) fused to a RT domain (M-MLV RT $^{D200N, L603W, T330P}$)

2) pegRNA
- sgRNA
- PAM sequence NGG
- gRNA scaffold
- Primer Binding Site (PBS)
- RT template including edit sequence

3) PE3 nicking sgRNA (to correct the non edited DNA strand)

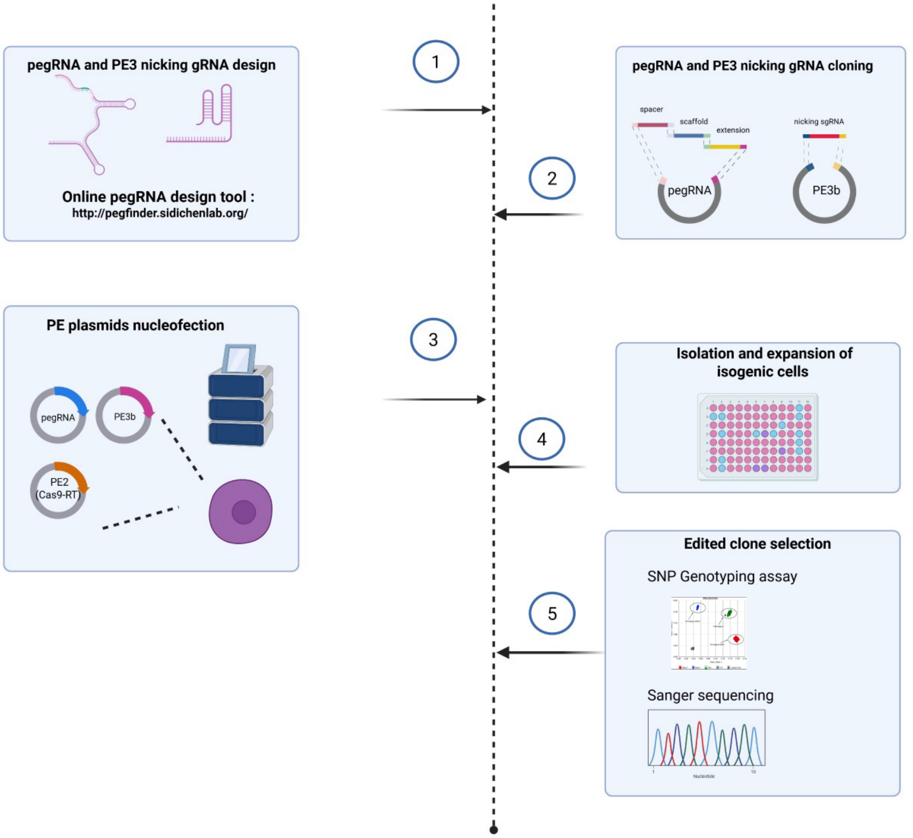


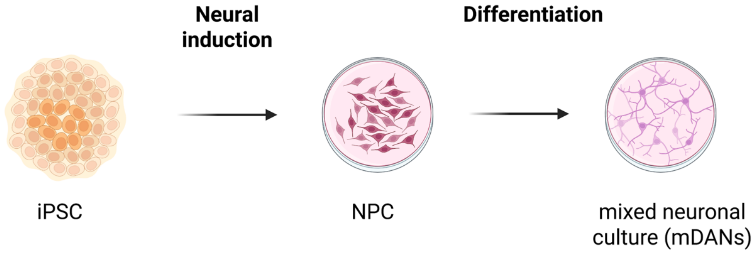

### Supplementary Figure 8

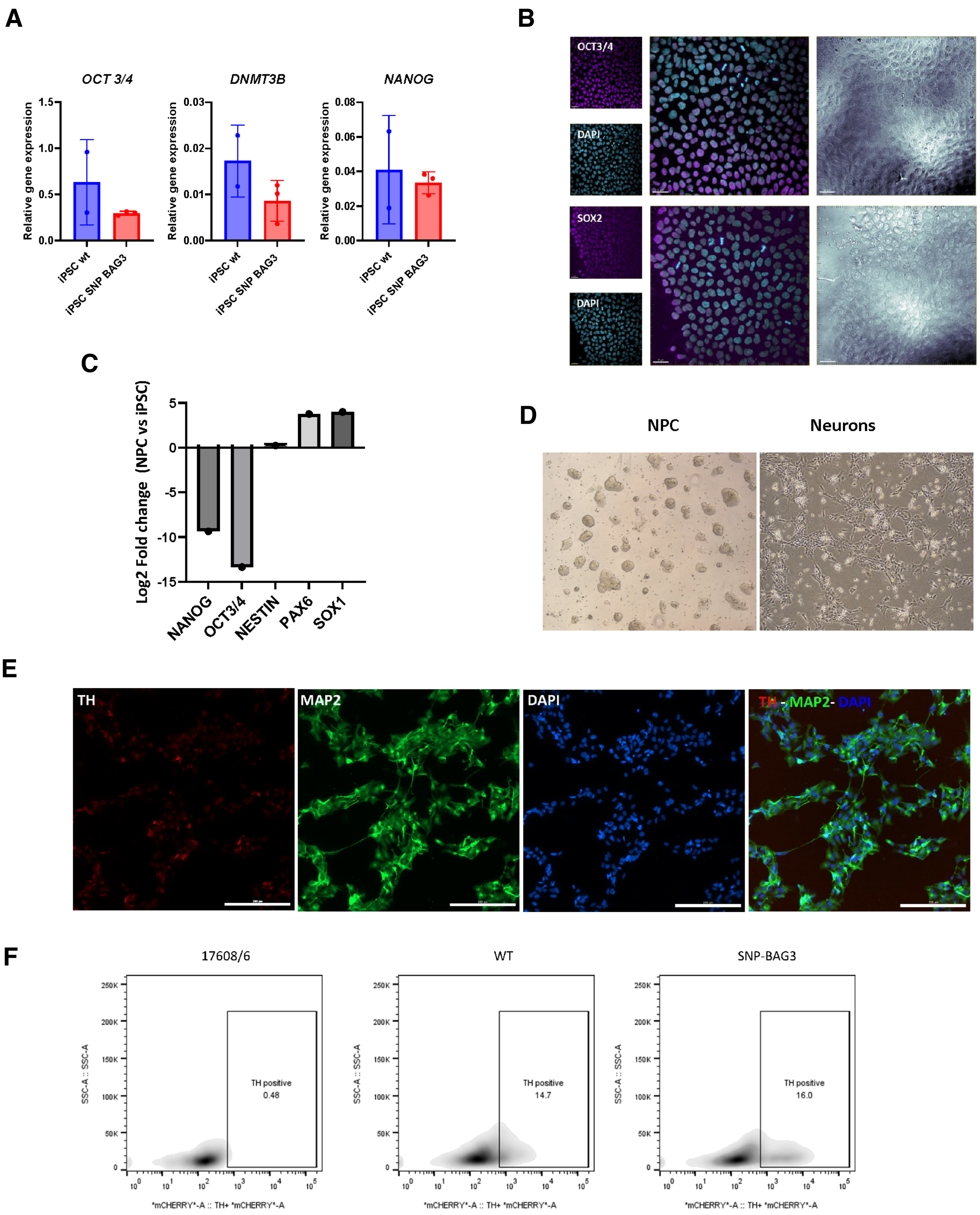

#### Supplementary Table legends

**Supplementary Table S1. Table of regulatory PD SNPs identified in smNPCs and mDANs using SNEEP.** The regulatory PD SNPs located in regulatory regions with potential to modify a TFBS in smNPCs and mDANs. The genomic coordinates, the rsID, GWAS p-value of variant association with PD, and minor allele frequency (MAF), the nucleotide of variant 1 (effect allele), the nucleotide of variant 2 (other allele), the affected TF motif(s), the strand(s) of the motif, the log-ratio of the binding scores of the two variants, and the p-value of the binding affinity change are indicated.

**Supplementary Table S2. Table of regulatory PD SNPs associated with correlated TF-target gene pairs.** PD SNPs predicted to be alter TF binding of TF and target genes showing absolute Pearson correlation above 0.5 are listed. The predicted TF, target gene, their Pearson correlation, p-value of correlation, the TF-target gene pair, p-value of binding energy change, expression level (RPKM) of both genes, and the genomic coordinates and rsID of the variant are shown.

**Supplementary Table S3. DNA-binding motifs of the HD-LIM family of TFs.** The JASPAR motif IDs, corresponding TF names, and motif logos of HD-LIM family TFs are shown.
